# Pericyte-to-endothelial cell signaling via vitronectin-integrin regulates blood-CNS barrier

**DOI:** 10.1101/2021.04.22.441019

**Authors:** Swathi Ayloo, Christopher Gallego Lazo, Shenghuan Sun, Wei Zhang, Bianxiao Cui, Chenghua Gu

## Abstract

Endothelial cells of blood vessels of the central nervous system (CNS) constitute blood-CNS barriers. Barrier properties are not intrinsic to these cells; rather they are induced and maintained by CNS microenvironment. Notably, the abluminal surface of CNS capillaries are ensheathed by pericytes and astrocytes. However, extrinsic factors from these perivascular cells that regulate barrier integrity are largely unknown. Here, we establish vitronectin, an extracellular-matrix protein secreted by CNS pericytes, as a regulator of blood-CNS barrier function via interactions with its integrin receptor, α5 in endothelial cells. Genetic ablation of vitronectin or mutating vitronectin to prevent integrin binding as well as endothelial-specific deletion of integrin α5 causes barrier leakage. Furthermore, vitronectin-integrin α5 signaling maintains barrier integrity by actively inhibiting transcytosis in endothelial cells. These results demonstrate that signaling from perivascular cells to endothelial cells via ligand-receptor interactions is a key mechanism to regulate barrier permeability.

## Introduction

The CNS requires an optimal and tightly regulated microenvironment for efficient synaptic transmission. This is achieved by blood-CNS barriers that regulate substance flux to maintain tissue homeostasis. Two such barriers are the blood-brain barrier (BBB) and blood-retina barrier (BRB) that are physiologically similar barriers separating the blood from brain and retina, respectively. The restrictive permeability of CNS endothelial cells that constitute these barriers is a result of specialized tight junctions and low rates of transcytosis, which limit substance exchange between blood and the CNS tissue (Andreone et al., 2017; Ben-Zvi et al., 2014; Chow and Gu, 2017; Langen et al., 2019; Reese and Karnovsky, 1967; Zhao et al., 2015).

Barrier properties are not intrinsic to CNS endothelial cells; they require active induction and maintenance from brain parenchyma cells (Armulik et al., 2010; Daneman et al., 2010; Heithoff et al., 2021; Stewart and Wiley, 1981). For example, Wnt ligands released by glia and neurons act on CNS endothelial cells to induce and maintain barrier properties (Daneman et al., 2009; Liebner et al., 2008; Stenman et al., 2008; Wang et al., 2012, 2019b). In contrast to glia and neurons, pericytes are directly in contact with capillary endothelial cells. Pericytes ensheathe capillary endothelial cells and share the same basement membrane with them allowing for elaborate cellcell signaling between these two cells. Intriguingly, the brain and retina have the highest pericyte to endothelial cell ratio compared to that in other tissues (Frank et al., 1987; Shepro and Morel, 1993). Indeed, mice with decreased pericyte coverage around endothelial cells exhibit leaky blood-CNS barriers (Armulik et al., 2010; Bell et al., 2010; Daneman et al., 2010; Park et al., 2017). However, how pericytes signal to endothelial cells to maintain barrier integrity is unknown.

Here, we identify vitronectin, a pericyte-secreted extracellular matrix (ECM) protein as an important regulator of barrier integrity. We establish that vitronectin regulates barrier function via binding to its integrin receptors on endothelial cells. Specifically, vitronectin is enriched in CNS pericytes, and mice lacking vitronectin as well as vitronectin mutant mice (*Vtn^RGE^*) that cannot bind integrin receptors exhibit barrier leakage. Moreover, we found that the RGD-ligand binding integrins, α5 and αv are expressed in CNS endothelial cells, and endothelial cell-specific acute deletion of α5 but not αv results in leaky barrier. We further demonstrate that barrier leakage observed in vitronectin mutant mice is due to increased transcytosis in CNS endothelial cells, but not due to deficits in tight junctions; and activation of integrin α5 by vitronectin inhibits endocytosis in CNS endothelial cells. Together, our results indicate that ligand-receptor interactions between pericyte-derived vitronectin and its integrin receptor expressed in CNS endothelial cells are critical for barrier integrity and may provide novel therapeutic opportunities for CNS drug delivery.

## Results

### Vitronectin is enriched in CNS pericytes compared to pericytes of peripheral tissues and its expression coincides with functional barrier formation

In order to identify pericyte candidate genes important for barrier function we took advantage of the retina as a model system. In the retinas of mice, CNS vessels invade the optic nerve head at postnatal day (P1) and expand radially from the center toward the periphery. As vessels grow from the proximal to distal ends of the retina, proximal vessels gain barrier properties and have a functional BRB, while the newly formed distal vessels have a leaky BRB (Chow and Gu, 2017). By examination of an existing transcriptome comparing the proximal and distal retinal vessels that contain a mixture of endothelial cells and pericytes (Strasser et al., 2010) and a brain pericyte transcriptomic database (He et al., 2016), we identified CNS pericyte genes that correlated with functional blood-retinal barrier. Of these candidate pericyte genes, we focused specifically on secreted and trans-membrane proteins as these are likely involved in ligand-receptor interactions. These analyses identified vitronectin, an extracellular matrix protein.

To validate our gene expression analyses, we first examined vitronectin protein localization in the CNS. Consistent with enriched *Vtn* transcript in proximal compared to distal vessels of the developing retina (Strasser et al., 2010), we also observed vitronectin protein highly expressed in proximal vessels with sealed BRB, and was not detectable in the distal, leaky vessels of the retina at P7 (Figures 1A, S1A). Similarly, vitronectin protein was also detected along capillaries of the brain (Figure 1B). As vitronectin is a secreted protein, to determine the cells producing *Vtn*, we used in situ hybridization to examine *Vtn* mRNA in the brain. *Vtn* mRNA is specifically expressed in *Pdgfrb* expressing pericytes adjacent to capillary endothelial cells (Figure 1C). At P7, *Vtn* is expressed in >97% (115 out of 118 cells across 3 mice) of the *Pdgfrb+* pericytes abutting capillaries and was present throughout the brain (Figures 1C, S1B). In contrast to high *Vtn* mRNA in CNS pericytes, we observed little to no expression in pericytes of peripheral tissues, such as the lung (Figures 1D, 1E). Our data are consistent with recent single-cell RNA sequencing studies revealing abundant *Vtn* expression in pericytes and little to no expression in non-vascular CNS cells such as neurons and astrocytes, both in the brain and the retina, through development and adulthood (Macosko et al., 2015; La Manno et al., 2021; Vanlandewijck et al., 2018).

**Figure 1.**
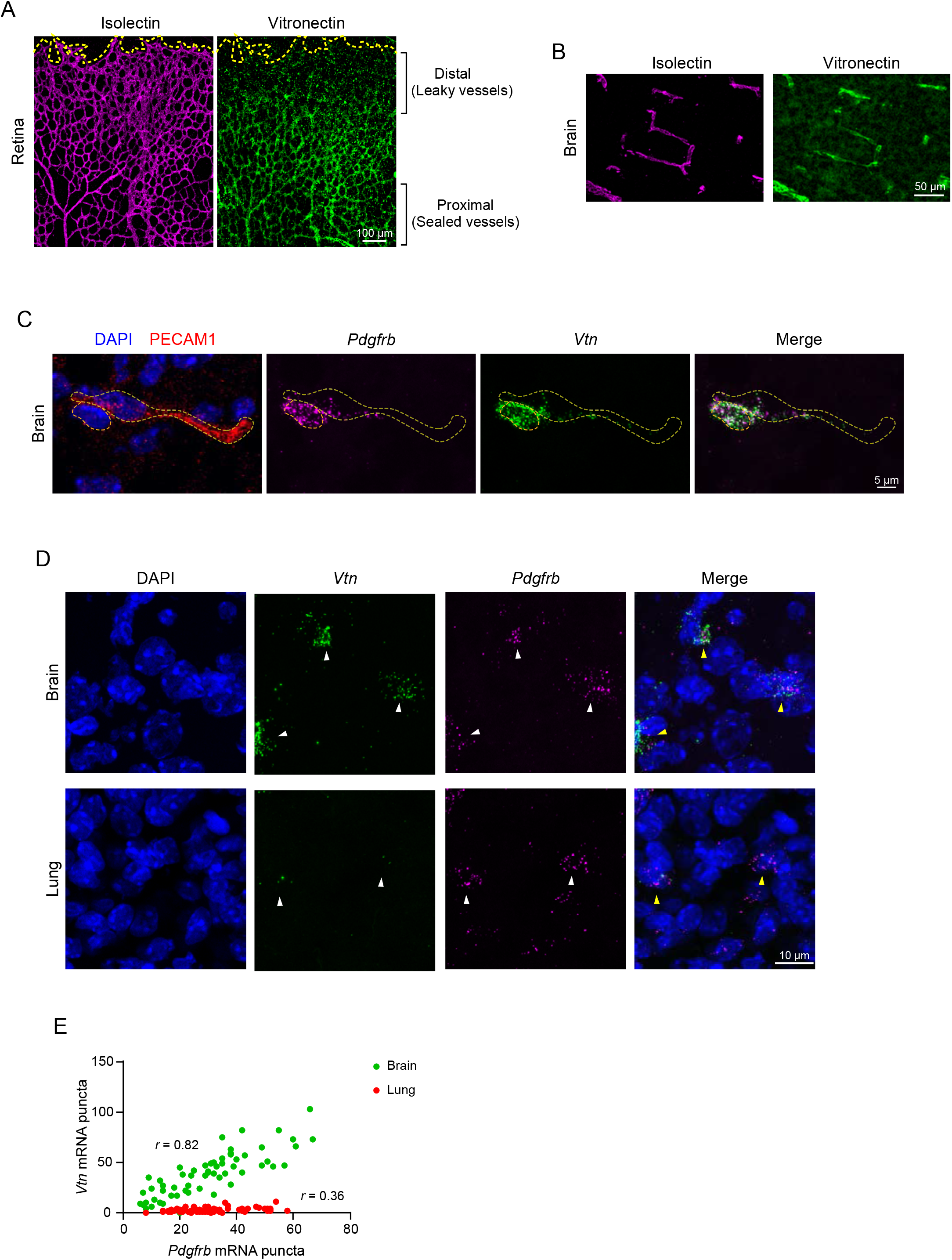
Vitronectin expression coincides with functional barrier formation and is enriched in CNS pericytes compared to peripheral tissue pericytes. (A, B) Immunostaining of vitronectin (green) and blood vessels (isolectin, magenta) in retina (A) and (B) brain of P7 mouse. (C) In situ hybridization for *Vtn* (green), *Pdgfrb* (pericyte gene, magenta) and immunostaining for PECAM1 (vessel marker, red) with DAPI (nuclei, blue) in P7 brain. Dashed lines indicate pericyte nucleus and vessel outline. (D) In situ hybridization for *Vtn* and *Pdgfrb* in brain and lung of P7 mouse. (E) Scatter plot between *Pdgfrb* and *Vtn* in brain (green) and lung (red). Each dot represents an individual cell, n=60 and 62 cells respectively from N=3 animals. Pearson correlation coefficient *r* = 0.82 for brain and *r* = 0.36 for lung.

### Pericyte-secreted vitronectin is required for blood-CNS barrier integrity

To determine if vitronectin is essential for barrier function, we performed a tracer leakage assay in P10 *Vtn^-/-^* mice when BRB and BBB are fully functional (Andreone et al., 2017; Chow and Gu, 2017). The injected tracer was completely confined to vessels in control mice, whereas tracer leaked out of vessels in the retinas of *Vtn^-/-^* mice. Numerous leaky hotspots of the tracer were apparent in the parenchyma (Figure 2A) and in neuronal cell bodies (Figure 2B) in the retinas of these mice (Figure 2C). This leakage was not limited to just small tracers like Sulfo-NHS-Biotin (0.5 kDa) as we observed similar leakage with higher molecular weight tracers such as 10 kDa Dextran (Figure S2A, S2B). Furthermore, this leakage persisted through adulthood in these mice (Figure S2C). Similar to the retina, we also observed BBB leakage in the cerebellum (Figures 2D, 2E). Thus, these findings demonstrate an important role for vitronectin in blood-CNS barrier function.

**Figure 2.**
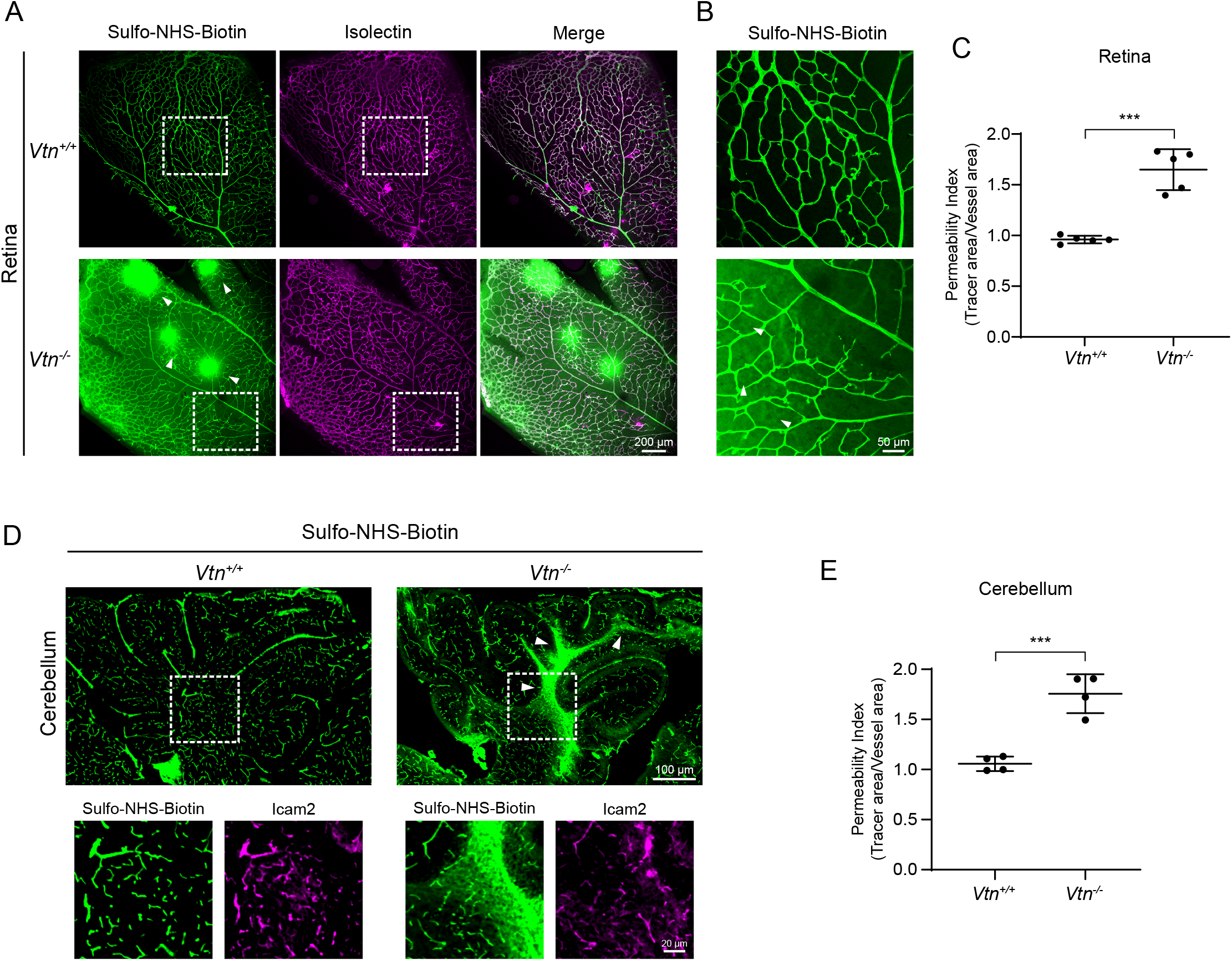
Vitronectin is essential for blood-CNS barrier integrity. (A) Sulfo-NHS-Biotin (tracer, green) leaks out of blood vessels (isolectin, magenta) in retinas of P10 *Vtn^-/-^* mice. Arrowheads show tracer hotspots in *Vtn^-/-^* mice, white boxes correspond to higher magnification images in (B). (B) Sulfo-NHS-Biotin leaked out in *Vtn^-/-^* mice taken up by neuronal cell bodies (arrowheads). (C) Quantification of vessel permeability in retinas of wildtype and *Vtn^-/-^* mice. n = 5 animals per genotype. Mean ± S.D.; ***p < 0.001; Student’s t-test. (D) Leakage of Sulfo-NHS-Biotin (green) from blood vessels (Icam2, magenta) in the cerebellum of P10 *Vtn^-/-^* mice. White boxes correspond to higher magnification panels shown. (E) Quantification of vessel permeability in retinas of wildtype and *Vtn^-/-^* mice. n = 4 animals per genotype. Mean ± S.D.; ***p < 0.001; Student’s t-test.

In addition to being an ECM protein, vitronectin is also an abundant protein circulating in the plasma which has been shown to mediate the complement pathway and play a role in tissue repair and wound healing (Leavesley et al., 2013; Preissner and Reuning, 2011). To fully establish that the barrier deficits we observe in *Vtn^-/-^* mice are indeed due to the lack of pericyte-secreted vitronectin and not due to the lack of circulating vitronectin, we took advantage of the fact that vitronectin in plasma is synthesized and secreted by liver hepatocytes and intravenous injections of siRNAs largely target the liver tissue with little to no distribution to other tissues (Song et al., 2003; Zender et al., 2003). To specifically knockdown the circulating vitronectin, we intravenously injected siRNAs targeting vitronectin or control siRNAs into 6-week old wildtype mice on two consecutive days (Figure 3A) based on previously established studies for efficient liver targeting (Song et al., 2003; Wrobel et al., 2021). Knockdown of vitronectin in plasma was evaluated by ELISA at day 3 or 5, i.e. 24 hours or 72 hours post the last dose of siRNA injections (Figure 3A). ELISA kit was validated by measuring levels of plasma vitronectin in wild-type, *Vtn^+/-^*, and *Vtn^-/-^* mice (Figure 3B). We performed these experiments with two independent siRNAs and at both the timepoints, for both the siRNAs, we observed >90% knockdown of vitronectin in plasma isolated from mice injected with vitronectin targeting siRNAs (Figures 3C, 3F). Importantly, at both these timepoints, we detected no leakage in mice injected with either siRNA targeting vitronectin or control siRNA (Figures 3D, 3E, 3G, 3H), demonstrating that plasma vitronectin is dispensable for barrier function. Thus, the combination of leaky barriers in *Vtn^-/-^* mice and intact barriers in mice specifically lacking plasma vitronectin demonstrates an important role for pericyte-secreted vitronectin in blood-CNS barrier function.

**Figure 3.**
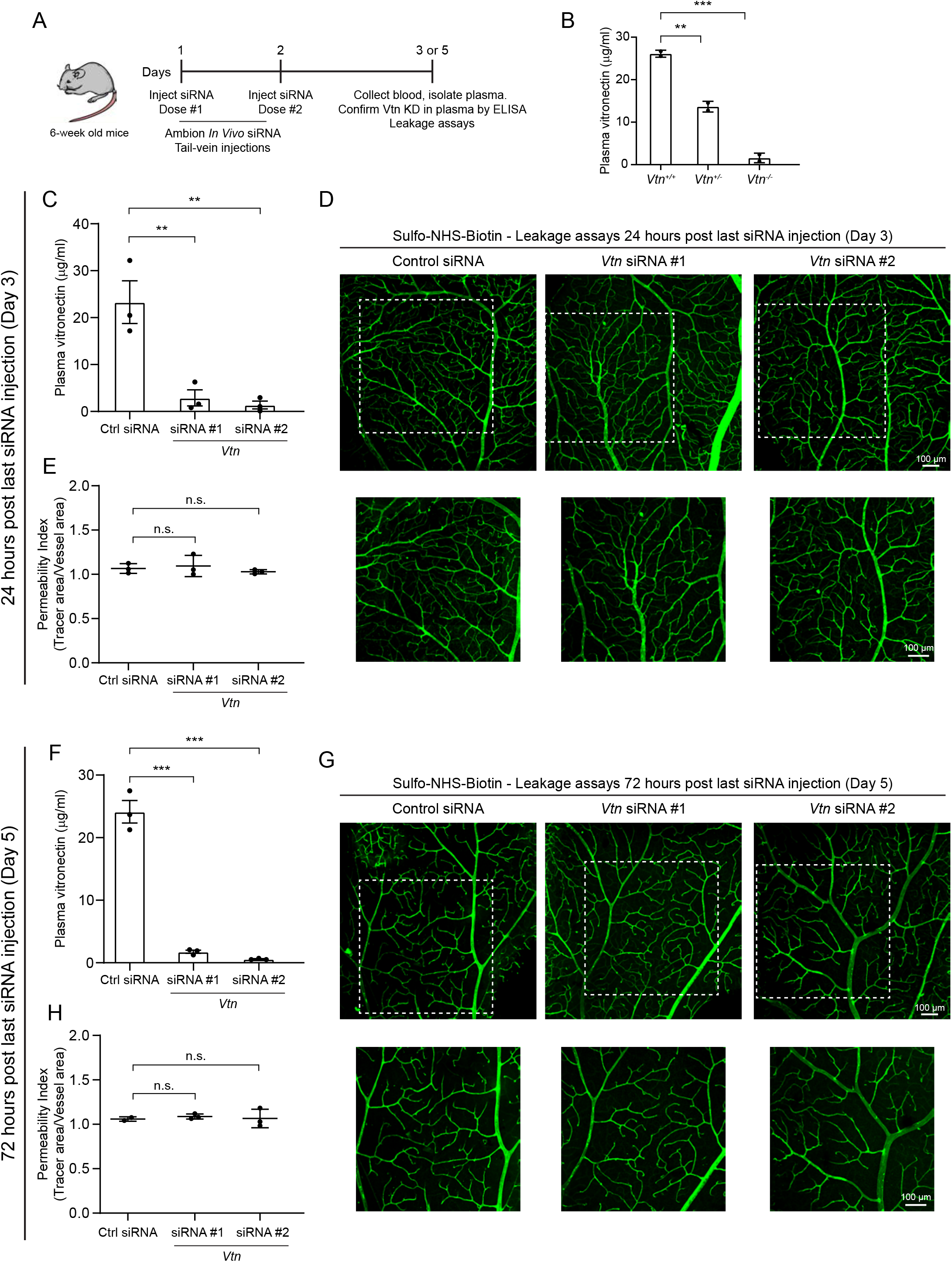
Vitronectin in plasma is not required for blood-CNS barrier function. (A) Illustration of experimental paradigm to knock-down plasma vitronectin specifically with intravenous injections of siRNAs followed by vitronectin protein level measurement in plasma and evaluation of barrier integrity by leakage assays. (B) Validation of ELISA kit to measure vitronectin protein levels in plasma of wildtype, heterozygotes and vitronectin null mice. n = 2 animals per genotype. Mean ± S.D.; **p < 0.01, ***p < 0.001; one-way ANOVA with Tukey’s post hoc test. (C, F) Vitronectin levels measured by ELISA in plasma of mice injected with control (Ctrl) siRNA or two independent siRNAs targeting vitronectin. 24 hours post last siRNA injection (C) and 72 hours post last siRNA injection (F). n = 3 mice per siRNA in each case. Mean ± S.D.; **p < 0.01, ***p < 0.001; one-way ANOVA with Tukey’s post hoc test. (D, E) Sulfo-NHS-Biotin confined to vessels (D) in retinas of mice injected with siRNA targeting vitronectin, 24 hours post last siRNA injection. Corresponding quantification of vessel permeability (E). White boxes correspond to panel of higher magnification images. n = 3 mice per siRNA. Mean ± S.D.; n.s. not significant, p > 0.05; one-way ANOVA with Tukey’s post hoc test. (G, H) Sulfo-NHS-Biotin confined to vessels (G) in retinas of mice injected with siRNA targeting vitronectin, 72 hours post last siRNA injection. Corresponding quantification of vessel permeability (H). White boxes correspond to panel of higher magnification images. n = 3 mice per siRNA. Mean ± S.D.; n.s. not significant, p > 0.05; one-way ANOVA with Tukey’s post hoc test.

### Vitronectin regulates blood-CNS barrier function by suppressing transcytosis in CNS endothelial cells

To determine the subcellular basis in endothelial cells for the underlying leakage we observed in *Vtn^-/-^* mice, we injected horse-radish peroxidase (HRP) intravenously and performed EM analysis in retina and cerebellum. In both retina and cerebellum of *Vtn^-/-^* mice and wildtype littermates, we observed HRP halting at tight junction “kissing points” between endothelial cells of capillaries (Figure 4A, 4B), indicating functional tight junctions. Consistent with this, we observed no changes in Claudin-5 protein expression in the retinas of *Vtn^-/-^* mice (Figure 4C). Closer examination of Claudin-5 and ZO-1 revealed that the localization of these proteins to cell-cell junctions is also unaltered (Figure 4D). In contrast, both retinal and cerebellum endothelial cells of *Vtn^-/-^* mice exhibited significantly increased HRP-filled vesicles (Figures 4E, 4G). Retinal endothelial cells had a 3-fold increase in vesicles compared to wildtype mice (Figure 4F) and cerebellar endothelial cells had 2-fold increase (Figure 4H), revealing that transcytosis is upregulated in these mice. Interestingly, in the cerebellum of these mice, we also observed HRP-filled vesicles in adjacent pericytes (Figure S3A) and about 25% of capillaries had their basement membrane completely filled with HRP (Figures S3B, S3C). This result indicates that tracer-filled vesicles are indeed transcytosed across endothelial cells (red arrowheads in example 2 of Figure S3A). These data reveal that barrier leakage in *Vtn^-/-^* mice is due to upregulated transcytosis. Together, these observations demonstrate that pericyte-derived vitronectin regulates barrier function by specifically inhibiting transcytosis in CNS endothelial cells.

**Figure 4.**
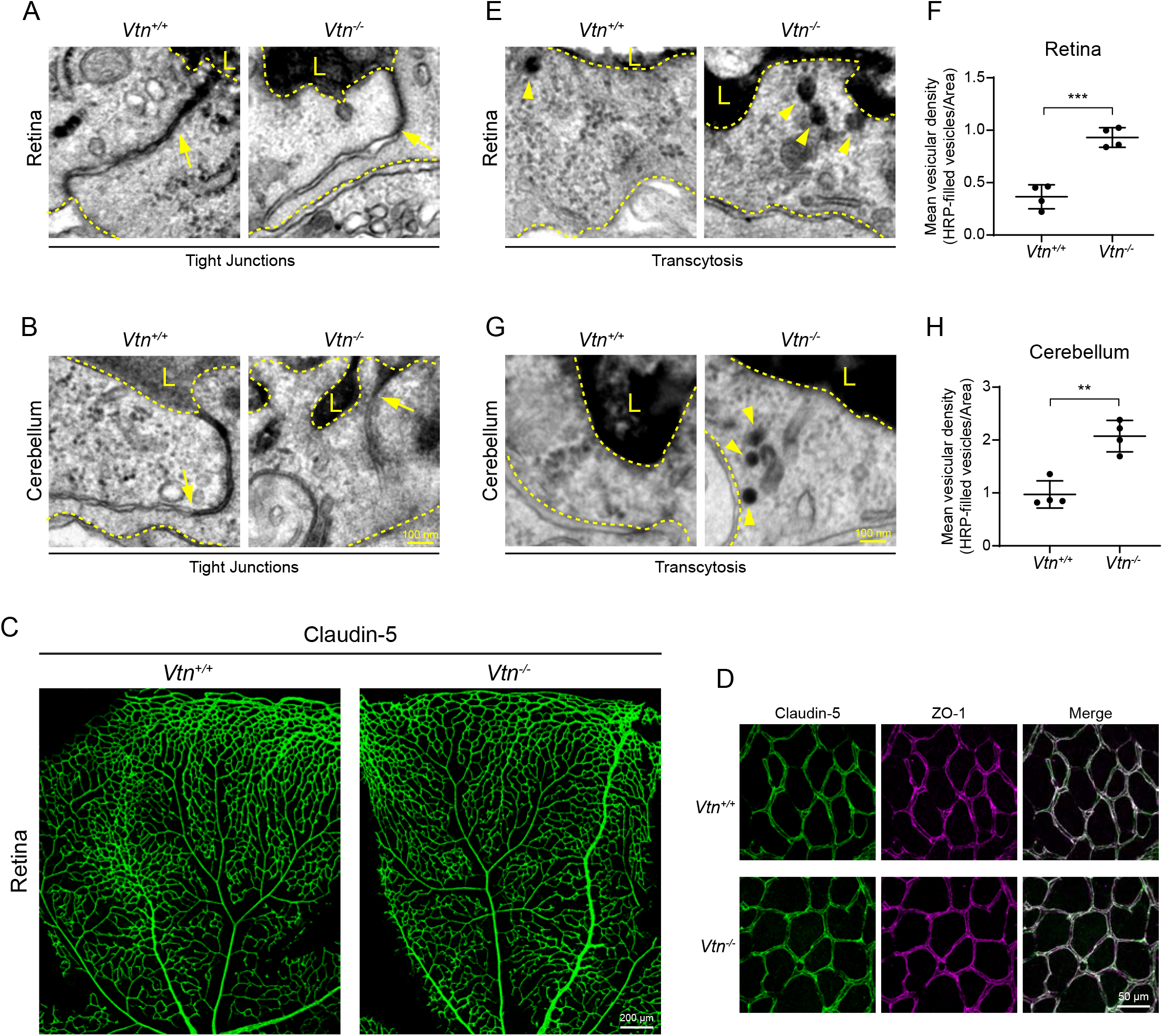
Vitronectin regulates blood-CNS barrier function by suppressing transcytosis in CNS endothelial cells. (A, B) EM images of HRP halting at tight junctions (arrows) in both retinas (A) and cerebellum (B) of wildtype and *Vtn^-/-^* mice. Luminal (L) and abluminal sides indicated by dashed yellow lines. (C) Claudin-5 immunostaining in P10 retinas of wildtype and *Vtn^-/-^* mice. (D) Higher magnification images of Claudin-5 (green) and ZO-1 (magenta) in P10 retinas to show expression and localization of tight junction proteins at cell-cell junctions. (E, G) EM images showing HRP-filled vesicles (arrowheads) in endothelial cells of retinas (E) and cerebellum (G) of wildtype and *Vtn^-/-^* mice. (F, H) Quantification of tracer-filled vesicles in endothelial cells of retinas (F) and cerebellum (H). n = 4 animals per genotype, 15-20 vessels per animal. Mean ± S.D.; ***p < 0.001, **p < 0.01; Student’s t-test.

### Vitronectin is not required for normal vessel patterning or pericyte coverage

How does vitronectin, a pericyte secreted protein regulate barrier properties in endothelial cells? One possibility is that barrier defects are due to impaired vessel development as pericytes are known to regulate endothelial sprouting and angiogenesis in the postnatal retinal vasculature (Eilken et al., 2017). However, we observed no changes in vascular patterning (Figure 5A). Specifically, we observed no changes in vessel density, capillary branching or radial outgrowth of the vasculature in *Vtn^-/-^* mice compared to wildtype littermates (Figures 5B-D).

**Figure 5.**
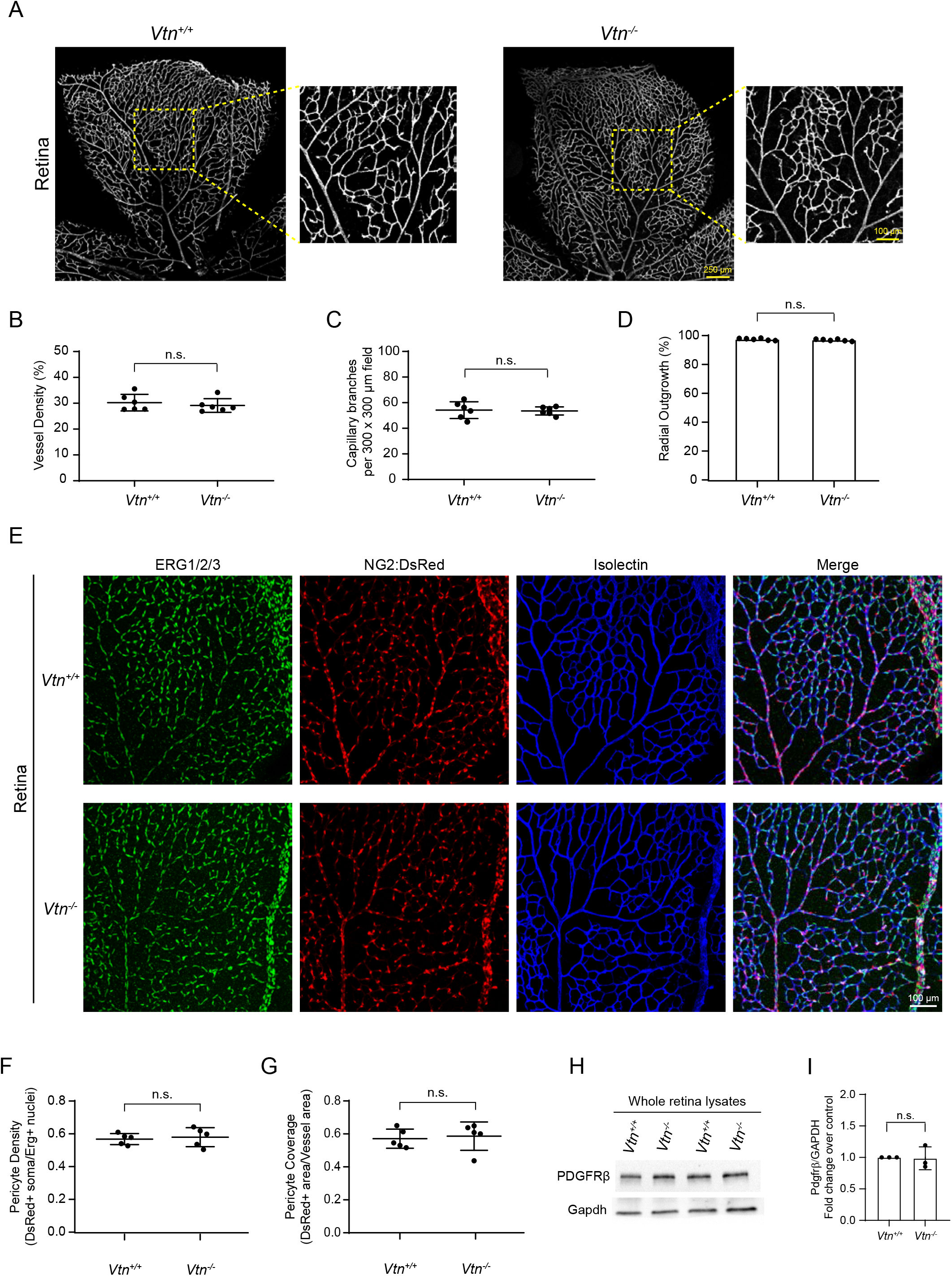
Vitronectin is not required for normal vessel patterning or pericyte coverage. (A) Tilescan images showing retinal vasculature of P10 retinas in wildtype and *Vtn^-/-^* mice. (B-D) Quantification of vessel density (B), capillary branching (C) and radial outgrowth (D) in P10 retinas. n = 6 animals per genotype. Mean ± S.D.; n.s. not significant, p > 0.05; Student’s t-test. (E) P10 retinas of NG2:DsRed+ wildtype and *Vtn^-/-^* mice immunostained for ERG1/2/3 (endothelial nuclei, green) and vessels (isolectin, blue). (F, G) Quantification of pericyte coverage (F) and pericyte density (G) in retinas of wildtype and *Vtn^-/-^* mice. n = 5 animals per genotype. Mean ± S.D.; n.s. not significant, p > 0.05; Student’s t-test. (H, I) Representative western blots (H) and quantification (I) of PDGFRβ protein levels normalized to Gapdh in whole retinal lysates of P10 wildtype and *Vtn^-/-^* mice. n = 3 animals per genotype. Mean ± S.D.; n.s. not significant, p > 0.05; Student’s t-test.

Another possibility is that since vitronectin is a pericyte gene and pericyte-deficient mice exhibit leaky barriers (Armulik et al., 2010; Daneman et al., 2010), barrier defects could be due to altered pericyte coverage. However, using NG2:DsRed mice that label mural cells (Figure 5E), similar pericyte coverage and pericyte density were observed in *Vtn^-/-^* and wildtype mice (Figures 5F, 5G). PDGFRβ protein levels were also similar in *Vtn^-/-^* and wildtype retinas (Figure 5H, 5I). Hence, vitronectin regulates barrier function without altering pericyte density or pericyte coverage.

Since vitronectin is an ECM protein, we also examined whether the barrier deficits in *Vtn^-/-^* mice were a consequence of structural and/or functional changes in the ECM. EM analysis of retinal and cerebellum capillaries revealed no obvious structural changes in the basement membrane of endothelial cells in *Vtn^-/-^* mice compared to wildtype mice. Moreover, immunostaining and western blotting of two highly enriched ECM proteins, collagen IV and fibronectin reveal normal localization (Figures S4A, S4B) and expression levels (Figure S4D, S4E) of these proteins in retinas of *Vtn^-/-^* mice. Notably, we also observed normal collagen IV ensheathment of retinal blood vessels in *Vtn^-/-^* mice (Figure S4C). Similarly, we observed normal localization and expression of two other ECM proteins, perlecan and laminin α4 in the retina vasculature of *Vtn^-/-^* mice (Figures S4F, S4G). Finally, we also used our EM data to investigate astrocyte endfeet attachments and observed no changes in average area of cerebellum capillaries covered by astrocyte endfeet between wildtype and *Vtn^-/-^* mice (Figure S4H, S4I). Thus, the overall structural and functional organization of the vascular basement membrane or the ECM was not compromised in mice lacking vitronectin.

Collectively, our data demonstrate that barrier defects caused in mice lacking vitronectin are not due to defects in vessel morphology and patterning, pericyte coverage, ECM organization or astrocyte endfeet. Rather, pericyte-secreted vitronectin could be acting directly on endothelial cells to regulate their barrier properties.

### Vitronectin binding to integrin receptors is essential for barrier function

How does pericyte secreted vitronectin signal to neighboring endothelial cells to regulate barrier permeability? Vitronectin belongs to the family of adhesion proteins that bind integrin receptors through a 3 amino acid motif, Arg – Gly – Asp (RGD) (Hynes, 2002; Preissner, 1991; Preissner and Reuning, 2011) (Figure 6A). Previous biochemical studies established that competitive binding experiments using short peptides containing the RGD domain (Orlando and Cheresh, 1991) as well as mutating vitronectin RGD to RGE (aspartic acid to glutamic acid) effectively abolished binding of vitronectin to integrins (Cherny et al., 1993). To determine if vitronectin binding to integrins is essential for barrier function, we took advantage of the knock-in mouse line harboring single point mutation of vitronectin with RGD domain mutated to RGE, *Vtn^RGE/RGE^* (Wheaton et al., 2016), hereafter referred to as *Vtn^RGE^*.

**Figure 6.**
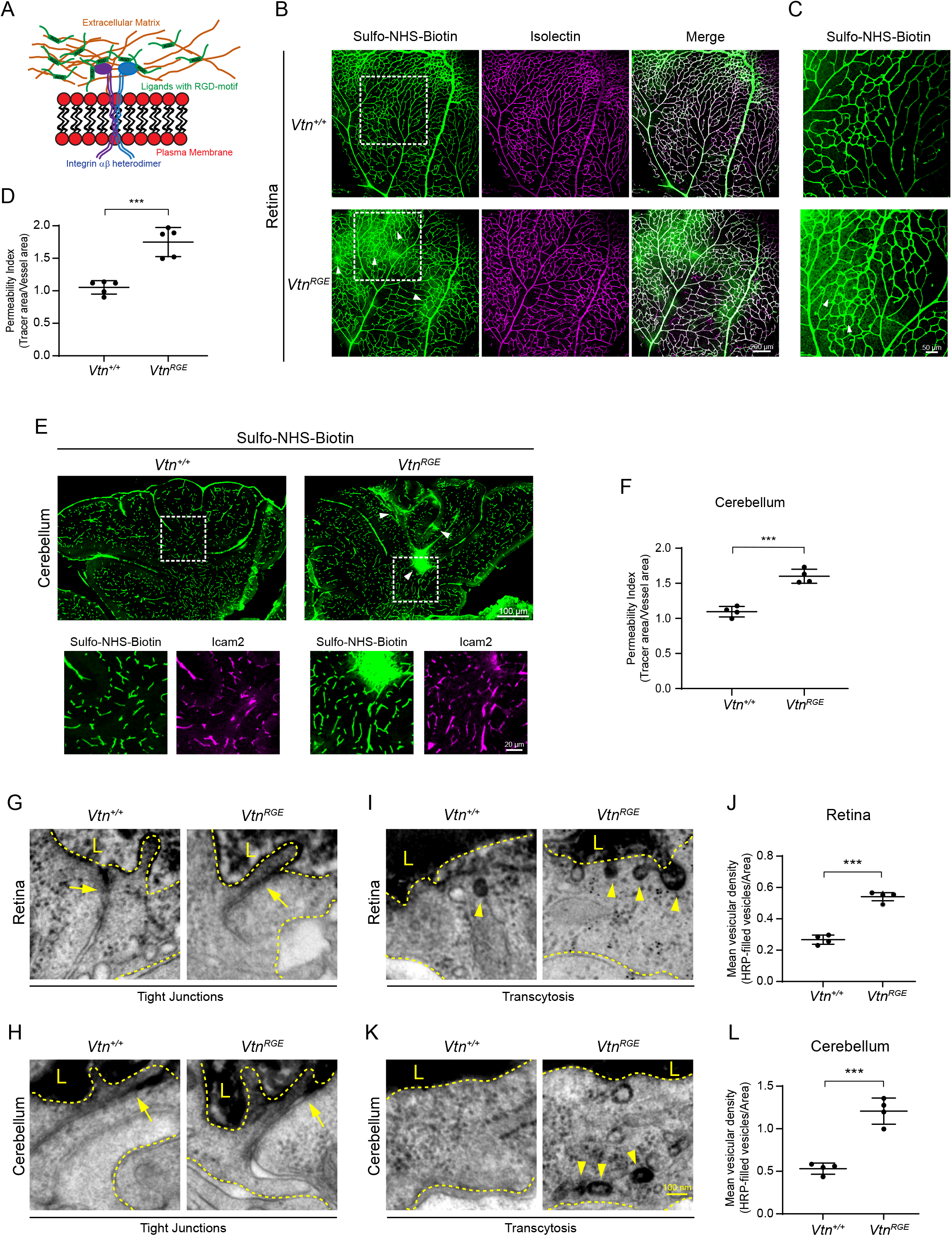
Vitronectin binding to integrin receptors is essential for barrier function. (A) Schematic illustrating binding of ligands containing RGD-motif with integrin receptors (B) Leakage of Sulfo-NHS-Biotin (tracer, green) from vessels (isolectin, magenta) in P10 *Vtn^RGE^* mice. White boxes correspond to higher magnification images shown in (C). (C) Tracer hotspots (arrowheads) in retinas of *Vtn^RGE^* mice. (D) Quantification of vessel permeability in wildtype and *Vtn^RGE^* mice. n = 5 animals per genotype. Mean ± S.D.; ***p < 0.001; Student’s t-test. (E) Leakage of Sulfo-NHS-Biotin (green) in the cerebellum of P10 *Vtn^RGE^* mice. White boxes correspond to higher magnification panels showing tracer confinement to vessels (ICAM2, magenta) in wildtype mice and tracer leakage in *Vtn^RGE^* mice. (F) Quantification of vessel permeability in wildtype and *Vtn^RGE^* mice. n = 4 animals per genotype. Mean ± S.D.; ***p < 0.001; Student’s t-test. (G, H) EM images of HRP halting at tight junctions (arrows) in both retinas (G) and cerebellum (H) of wildtype and *Vtn^RGE^* mice. Luminal (L) and abluminal sides indicated by dashed yellow lines. (I, K) EM images of HRP-filled vesicles (arrowheads) in endothelial cells of retinas (I) and cerebellum (K) of wildtype and *Vtn^RGE^* mice. (J, L) Quantification of tracer-filled vesicles in endothelial cells of retinas (J) and cerebellum (L). n = 4 animals per genotype, 15-20 vessels per animal. Mean ± S.D.; ***p < 0.001, **p < 0.01; Student’s t-test.

*Vtn^RGE^* mice exhibited leakage in retina and cerebellum, phenocopying *Vtn^-/-^* mice. Similar hotspots of tracer leakage (Figure 6B) as well as neuronal cell bodies filled with tracer were apparent in the retinal parenchyma of *Vtn^RGE^* mice (Figures 6C, 6D) with no obvious defects in vascular patterning. The BBB in the cerebellum of these mice was also leaky with tracer hotspots throughout the cerebellum (Figures 6E, 6F).

EM analysis of retinas and cerebellum of HRP-injected *Vtn^RGE^* mice revealed similar subcellular features in endothelial cells as observed in the *Vtn^-/-^* full knockout mice. Both the retina and cerebellum in *Vtn^RGE^* mice had functional tight junctions as noted by HRP halting sharply between endothelial cells (Figures 6G, 6H). However, we observed >2-fold increase in HRP-filled vesicles in endothelial cells of *Vtn^RGE^* mice compared to wildtype animals (Figures 6I, 6J, 6K, 6L), revealing upregulated transcytosis in these mice. Thus, *Vtn^RGE^* mice fully recapitulate the phenotype observed in *Vtn^-/-^* mice, indicating vitronectin binding to integrin receptors is critical for its role in regulating barrier function.

### Vitronectin-integrin α5 signaling inhibits endocytosis in CNS endothelial cells

What are the specific integrin receptors on endothelial cells required for vitronectin-mediated regulation of barrier function? Of the several α and β integrin subunits that heterodimerize to form integrin receptors (Hynes, 2002) we focused on the two α receptors, α5 and αv that have been shown to interact with RGD-ligands. While αv is considered the predominant receptor for vitronectin, cell motility studies have also demonstrated interaction of vitronectin with α5 (Bauer et al., 1992; Zhang et al., 1993). Indeed, when primary mouse brain endothelial cells were grown on vitronectin-coated dishes, we readily observed adhesive structures containing endogenous integrin α5 (Figures S5A). In contrast, no adhesion structures were observed in cells grown on collagen IV coated or laminin-511 (α5β1γ1, hereafter referred to as laminin) coated dishes, demonstrating specific binding and activation of integrin α5 by vitronectin (Figures S5A, S5B). To mimic and recapitulate the *in vivo* ECM as much as possible we also grew these cells on a combination of ECM ligands. Similar to our findings with individual ligands, we observed integrin α5 containing adhesion structures only in the presence of vitronectin (Figures 7A, 7B). Furthermore, >95% of α5 adhesions were positive for and co-localized with classic focal adhesion proteins such as phosphorylated FAK (Y397), paxillin, and vinculin (Figures S5C, S5D), indicating that α5 adhesions are indeed bonafide focal adhesion structures. Thus, vitronectin actively engages integrin α5 receptors in CNS endothelial cells and drives the formation of α5 containing focal adhesions.

Since deletion of vitronectin results in barrier leakage due to increased transcytosis, we next directly investigated the role of vitronectin-integrin α5 interactions in vesicular trafficking in primary brain endothelial cells. Knockdown of integrin α5 in these cells with two independent shRNAs (Figure 7C, 7D), resulted in a significant increase in endocytosis (Figures 7E, 7F). In cells depleted of integrin α5 by either shRNA, FM1-43FX dye was readily incorporated in the newly formed vesicles that were endocytosed from plasma membrane, whereas very few FM1-43FX dye-positive vesicles were observed within the cells transfected with scrambled shRNA. These results demonstrate that the engagement of vitronectin-integrin α5 actively inhibits endocytosis in CNS endothelial cells.

**Figure 7.**
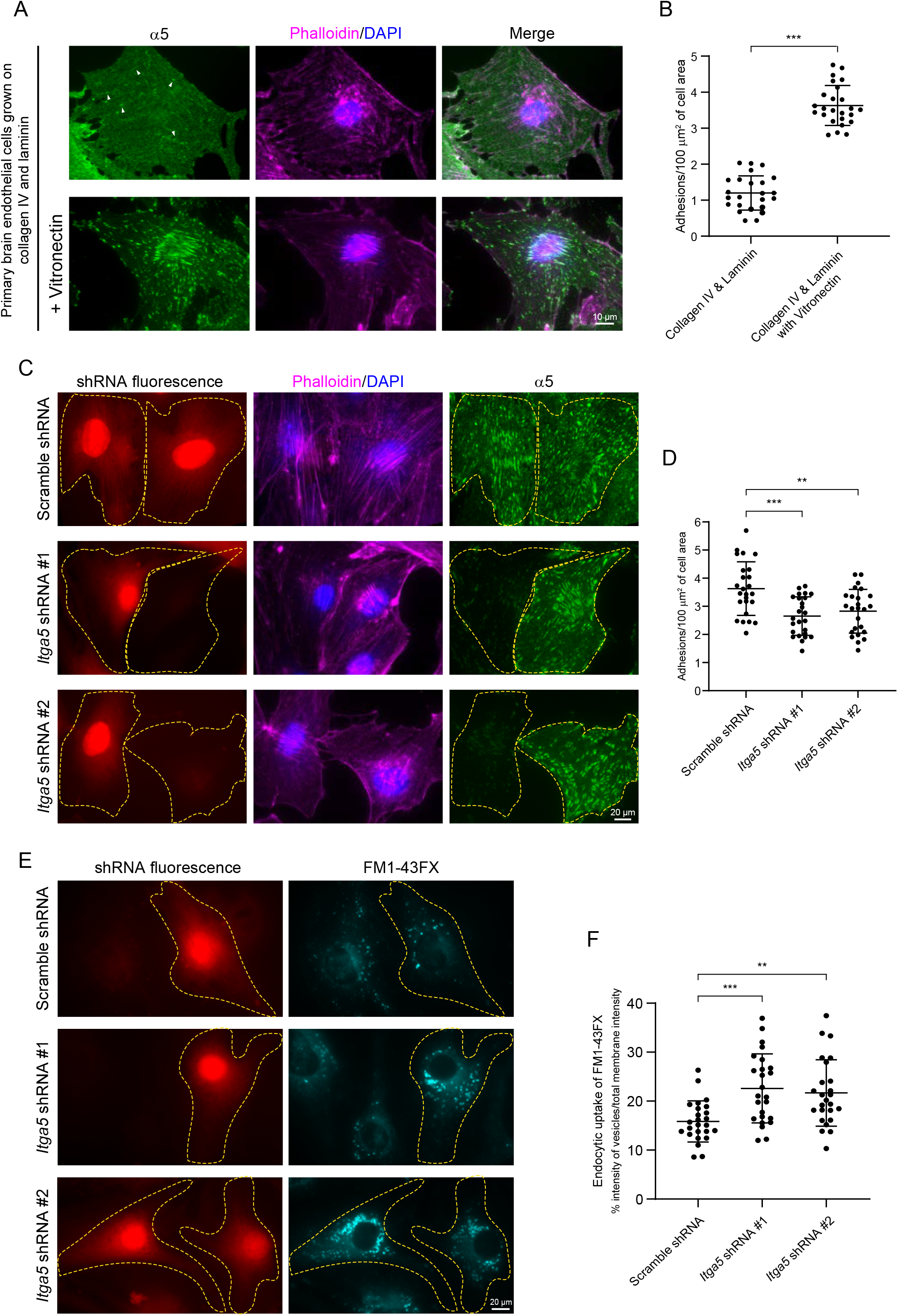
Engagement of integrin α5 with vitronectin forms adhesion structures and actively inhibits endocytosis in primary brain endothelial cells. (A, B) Representative images (A) and quantification (B) of α5 (green) containing adhesion structures in primary brain endothelial cells (phalloidin in magenta) grown on collagen IV, laminin and vitronectin-coated dishes. n = 25 cells per condition from 3 independent experiments. Mean ± S.D.; ***p < 0.001; Student’s t-test (C, D) Validation of two independent shRNAs (red) targeting endogenous α5 (green) in primary brain endothelial cells (phallodin in magenta) and quantification of adhesion structures (D) in scramble vs Itga5 shRNAs. n = 25 cells per condition from 3 independent experiments. Mean ± S.D.; ***p < 0.001, **p < 0.01; one-way ANOVA with Tukey’s post hoc test. (E) Endocytosis assay with membrane impermeable FM1-43FX (cyan) in primary brain endothelial cells transfected with shRNAs (red) targeting endogenous α5. (F) Quantitation of endocytic uptake of FM1-43FX. n = 25 cells per condition from 3 independent experiments. Mean ± S.D.; ***p < 0.001, **p < 0.01; one-way ANOVA with Tukey’s post hoc test.

### Integrin receptor, α5, is required in endothelial cells for blood-CNS barrier integrity

To identify the relevant endothelial integrin receptors *in vivo,* we first performed *in situ* hybridization to examine the localization of *Itga5* and *Itgav* transcripts in the brain. Single cell RNA-seq data reveals transcripts for both genes in CNS endothelial cells (Vanlandewijck et al., 2018); α5 and αv are also known to co-operate during vascular development (Van Der Flier et al., 2010). RNAscope reveals that although both *Itga5* and *Itgav* mRNA are present in endothelial cells as well as pericytes in the brain (Figure 8A, S7A), *Itga5* mRNA is predominantly expressed in endothelial cells whereas *Itgav* mRNA is predominately expressed in pericytes (Figure 8B). On average, *Itga5* transcripts in endothelial cells was 20.3 ± 1.5 and in pericytes it was 5.3 ± 0.43. In contrast, the average *Itgav* transcripts in endothelial cells was 8.3 ± 0.7 and in pericytes it was 19.5 ± 1.2 (Figure 8B, Mean ± SEM). Thus, *Itga5* is predominantly expressed in endothelial cells while *Itgav* is predominantly in perciytes.

**Figure 8.**
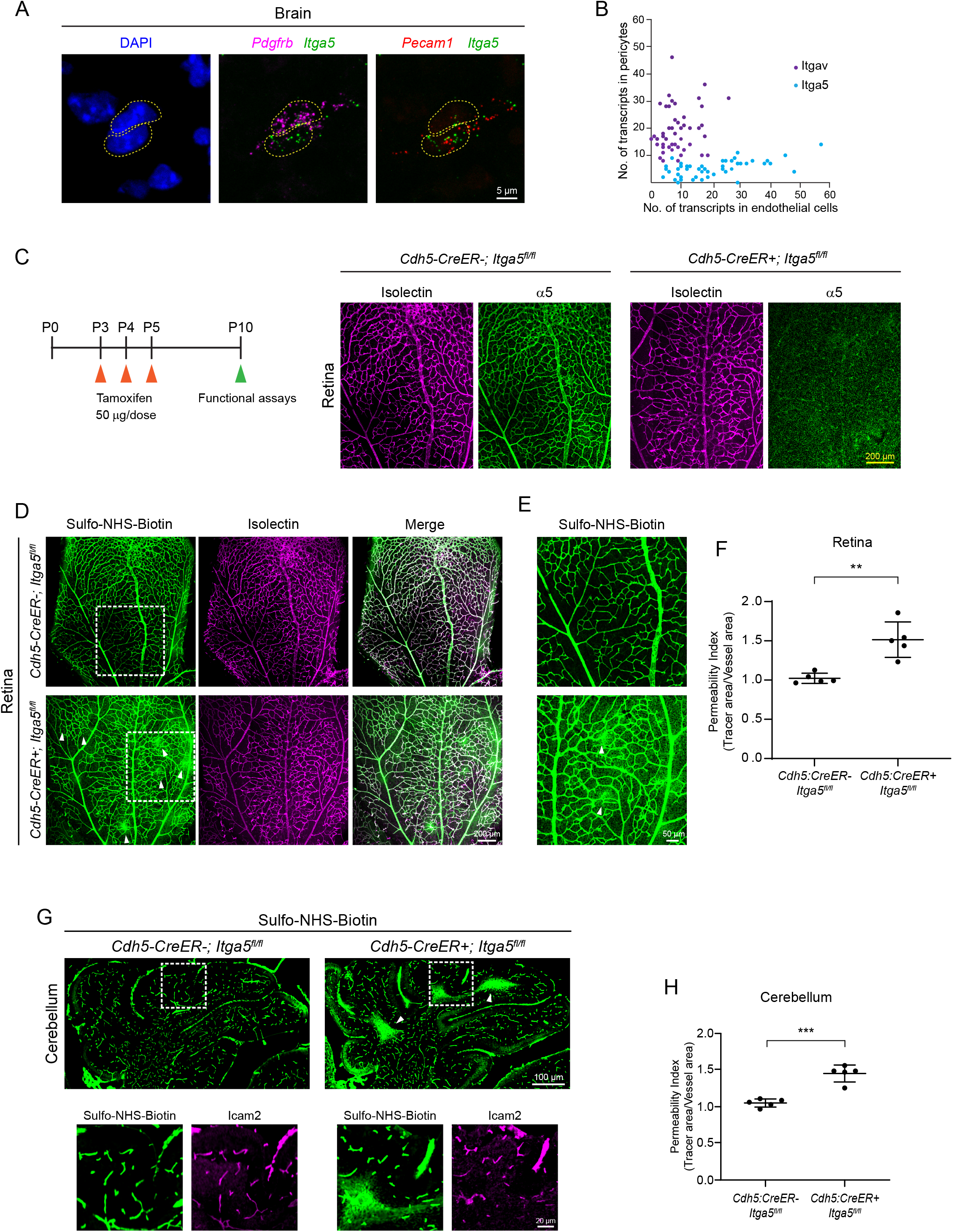
Integrin α5 in endothelial cells is specifically required for blood-CNS barrier function. (A) In situ hybridization for *Itga5* (green), *Pecam1* (endothelial gene, red) and *Pdgfrb* (pericyte gene, magenta) in P7 brain tissue. (B) Scatter plot of *Itga5* (blue) and *Itgav* (purple) transcript numbers in pericytes versus endothelial cells from RNAscope in situ hybridization as shown in (A). Each dot represents an individual pericyte-endothelial cell pair, n = 49 and 47 cell pairs respectively from 3 animals. (C) Depiction of tamoxifen injections in postnatal pups from P3-P5 and validation of tamoxifen-induced deletion of α5 (green) from vessels (isolectin, magenta) in P10 mice. (D) Sulfo-NHS-Biotin (tracer, green) leaks out of vessels (isolectin, magenta) in P10 *Cdh5:CreER+; Itga5^fl/fl^* mice. Tamoxifen administered P3-P5, see Figure S6. White boxes correspond to higher magnification images in (E). (E) Tracer hotspots (arrowheads) in mice lacking endothelial Itga5. (F) Quantification of vessel permeability in wildtype and *Cdh5:CreER+; Itga5^fl/fl^* mice. n = 5 animals per genotype. Mean ± S.D.; **p < 0.01; Student’s t-test. (G) Sulfo-NHS-Biotin (green) leakage in cerebellum of P10 *Cdh5:CreER+; Itga5^fl/fl^* mice. White boxes correspond to higher magnification images with tracer and vessels (ICAM2, magenta). (H) Quantification of vessel permeability in wildtype and *Cdh5:CreER+; Itga5^fl/fl^* mice. n = 5 animals per genotype. Mean ± S.D.; ***p < 0.001; Student’s t-test.

We next determined the role of endothelial *Itga5* and *Itgav* in barrier function by ablating these genes acutely and specifically in endothelial cells by crossing *Itga5^fl/fl^* or *Itgav^fl/fl^* (Van Der Flier et al., 2010) mice with endothelial cell-specific *Cdh5-CreER* mouse line (Wang et al., 2010). Acute deletion of endothelial *Itga5* (Figure 8C) resulted in a leaky blood-retinal barrier (Figure 8D) with numerous tracer hotspots spread out in the retina tissue (Figures 8E, 8F). Importantly, *Itga5^fl/fl^;Cdh5-CreER* mutant mice exhibited normal vascular density and patterning (Figure S6A-S6D). Similar to *Vtn^-/-^* and *Vtn^RGE^ mice, Itga5^fl/fl^;Cdh5-CreER* mutants also had a leaky BBB in the cerebellum (Figures 8G, 8H). In contrast to *Itga5*, mice lacking endothelial *Itgav* exhibited normal barrier function. The injected tracer was completely confined to the vasculature in both the retina (Figures S7B-S7D) and the brain tissue (Figures S7E, S7F) indicating intact blood-CNS barriers in *Itgav^fl/fl^;Cdh5-CreER* mice. These experiments establish the RGD-specific integrin receptor, α5 as an essential integrin for barrier function, particularly in the retina and cerebellum. Importantly, mice lacking endothelial integrin α5 receptor fully phenocopy *Vtn^-/-^* and *Vtn^RGE^* mice. Together with our *in vitro* results, these data indicate that the engagement of vitronectin-integrin α5 actively inhibits endocytosis in CNS endothelial cells to ensure barrier integrity. These results demonstrate that ligand-receptor interactions between pericyte-derived vitronectin and endothelial integrin, α5, are critical for barrier integrity by actively inhibiting transcytosis in CNS endothelial cells *in vivo*.

## Discussion

In this study, we establish how pericytes signal to endothelial cells via vitronectin-integrin interactions to maintain low rates of transcytosis, thus, ensuring barrier integrity. We identify and demonstrate an essential role for pericyte-secreted vitronectin in blood-CNS barrier function. Vitronectin deposited in the extracellular matrix, binds to integrin receptors on CNS endothelial cells and this interaction suppresses transcytosis, thus, regulating barrier integrity.

Lack of vitronectin or mutating vitronectin to prevent integrin binding as well as endothelial-specific integrin deletion – all of these genetic manipulations result in dysfunctional blood-CNS barriers, highlighting the role of ligand-receptor interactions of pericyte-to-endothelial signaling in barrier function.

How does pericyte-derived vitronectin interacting with integrin receptors actively inhibit endocytosis in CNS endothelial cells? The adhesive forces generated by the engagement of integrins have been shown in various cell types to maintain plasma membrane tension (Katsumi et al., 2004; Ross et al., 2013). Disengaging integrin receptors from vitronectin likely causes a reduction in membrane tension and it is well known that decreased membrane tension promotes increased endocytosis (Diz-Muñoz et al., 2013; Sheetz and Dai, 1996; Thottacherry et al., 2018). Thus, it is plausible that vitronectin binding to integrin α5 exerts adhesive forces to maintain the plasma membrane tension to ensure low rates of transcytosis in CNS endothelial cells. Our results indicate that signals from perivascular cells to endothelial cells via designated ligand-receptor pairs contribute to the unique biophysical properties of CNS endothelial membrane for barrier integrity.

Our work demonstrates the importance of signaling between perivascular cells and CNS endothelial cells in modulating blood-CNS barriers. It is likely that in addition to vitronectin, other ECM molecules also play key roles in barrier function given that ECM is a hub for ligand-receptor interactions across multiple cell types. Indeed, there is evidence for fibronectin interacting with α5 receptor impacting brain endothelial cell survival (Wang and Milner, 2006) and promoting blood-brain barrier integrity following stroke (Wang et al., 2019a). Absence of integrin receptors also triggers onset of experimental autoimmune encephalomyelitis (Kant et al., 2019). Not surprisingly, ECM breakdown is a hallmark feature associated with many diseases and disorders of the CNS (Thomsen et al., 2017). Thus, our findings reveal new molecular targets and pathways within the CNS for development of novel therapeutics that could aid CNS drug delivery.

## Acknowledgements

The authors thank all members of the Gu laboratory, particularly Hannah Zucker for comments on this manuscript; special thanks to Shachar Dagan, Sarah Pfau and Joseph Amick for technical assistance with *in vivo* siRNA experiments; Thomas Sisson for providing the *Vtn^-/-^* and *Vtn^RGE^* mice, Ralf Adams for *Cdh5:CreER* mice.

## Author contributions

S.A. and C.G. conceived the project and designed experiments. S.A., C.G.L. and S.S. performed experiments and analyzed all data. W.Z and B.C designed *in vitro* cell culture experiments. W.Z performed and analyzed cell culture experiments. S.A. and C.G. wrote the manuscript with all other authors’ input.

## Data and materials availability

Requests for data supporting figures and materials should be addressed to C.G.

