## Supplementary Material for "Pericyte-to-endothelial cell signaling via vitronectin-integrin regulates blood-CNS barrier"

This PDF file includes:

Materials and methods

Figures S1-S7

### Material and methods

#### Mice

All mouse experiments were performed according to institutional and US National Institutes of Health (NIH) guidelines approved by the International Animal Care and Use Committee (IACUC) at Harvard Medical School. Mice were maintained on 12 light/12 dark cycle. The following mice strains were used in this study: Both *Vtn* null mice (JAX: 004371) and $Vtn^{RGE/RGE}$ mice (Wheaton et al., 2016) were generous gifts from Dr. Thomas Sisson, University of Michigan; *Cdh5-CreER* was kindly provided by Dr. Ralf Adams, Max-Planck Institute of Molecular Biomedicine; *NG2:DsRed* (JAX: 008241), $Itga5^{flox}$ (JAX: 032299), $Itgav^{flox}$ (JAX: 032297); were obtained from Jackson Laboratories. All mice were maintained on C57/BL6J background.

Cre-mediated recombination was induced by intraperitoneal injection of 50 ug of 1 mg/ml Tamoxifen (T5468, Sigma dissolved in peanut oil containing ethanol 1:40 by volume) for 3 consecutive days, P3-P5. All animals were genotyped with allele-specific PCR reactions prior to experiments and both males and females were used for all experiments.

#### Immunohistochemistry

Enucleated eyes of P10 pups were fixed in 4% ice-cold PFA (EMS, 15713) for 5 minutes at room temperature (RT). Retinas were then dissected in 4% PFA and allowed to fix for 30 minutes at RT (adult retinas were fixed for 1 hour at RT). Retinas were washed in PBS three times for 5 minutes each and incubated in blocking buffer (10% normal donkey serum with 5% bovine serum albumin and 0.5% triton in PBS) for 1 hour at RT. Retinas were then incubated with primary antibodies in blocking buffer overnight at 4°C. The next day retinas were washed again with PBS three times for 5 minutes each and incubated with secondary antibodies in blocking buffer for 1 hour at RT and washed again with PBS. Retinas were then dissected into 4 leaflets and flat mounted on glass slides with vitreal surface in contact with coverslips using Prolong gold Antifade mountant (Thermo Fisher Scientific P36934)

Dissected brains were drop-fixed in 4% PFA overnight at 4°C. The brains were washed with PBS three times 10 minutes each and cryopreserved in 30% sucrose. The brains were bisected along the midline to obtain sagittal sections, before freezing in TissueTek OCT (Sakura). 20 um cryosections obtained on the cryostat were then processed for immunostaining. Brain sections were first permeabilized with 0.2% triton in blocking buffer (10% normal donkey serum with 5% bovine serum albumin in PBS) for 10 minutes at RT. The rest of the staining procedure was similar to the stainings in retina as above, except secondary antibodies were incubated for 45 minutes at RT.

Isolectin GS-$IB_4$ conjugated to Alexa Fluor 568 (Thermo Fisher Scientific I21412) or Isolectin GS-$IB_4$ conjugated to Alexa Fluor 647 (Thermo Fisher Scientific I32450) was incubated (1:300) along with primary antibodies to stain vasculature in retinas. Streptavidin conjugated to Alexa Fluor 647 (Thermo Fisher Scientific S32357) was incubated (1:300) along with secondary antibodies to detect the injected Sulfo-NHS-Biotin. The following primary antibodies were used: rabbit α-Vitronectin (Genway Biotech GWB-794F8F, 1:100), goat α -CD31 (PECAM1, R&D Systems AF3628, 1:100), mouse α-Claudin-5 (Thermo Fisher Scientific 352588, 1:200), rabbit α-ZO-1 (Invitrogen 40-2200, 1:200), rat α-CD102 (ICAM2, BD Biosciences 553326, 1:100), rabbit α-ERG1/2/3 (Abcam ab92513, 1:200), rabbit α-Collagen IV (Bio-rad 2150-1470, 1:200), rat α-Perlecan (EMD Millipore, MAB1948P, 1:200), goat α-laminin α4 (R&D systems, AF3837, 1:100) rat α-CD49e (α5, BD Biosciences 553319, 1:100). All corresponding secondary antibodies were used at 1:300 obtained from Jackson Immunoresearch Laboratories.

**Fluorescent in situ hybridization**

Brains and lungs were dissected from 1-week old wildtype animals, flash frozen in liquid nitrogen and cryosectioned to obtain 20 um sections. RNAscope was performed according to manufacturer's instructions (ACD Bio) with the following probes: *Vtn* (443601), *Pdgfrb* (411381-C3), *PECAM1* (316721-C2), *Itga5* (575741) *Itgav* (513901). To perform immunostaining post RNAscope, slides were briefly rinsed in PBS and incubated with blocking buffer (3% normal donkey serum with 3% bovine serum albumin in PBS) for 1 hour at RT. The rest of the steps for staining are similar to the staining protocol mentioned above under immunohistochemistry. DAPI (Thermo Fisher Scientific 62247) was added at 1:5000 dilution in the last PBS wash before mounting slides.

**Barrier permeability assays**

P10 pups were briefly anaesthetized with 3% isoflurane. Eyelid of one eye was cut off and tracer was injected retro-orbitally (Yardeni et al., 2011) with a 30-gauge needle. Tracer was allowed to circulate for 5 minutes followed by dissection of contralateral retina and brain, and processed for immunohistochemistry.

Tracers include EZ-Link Sulfo-NHS-LC-Biotin (Thermo Fisher Scientific, 21335, 0.44 kDa) injected at 0.5 mg/gm body weight and 10 kDa Dextran conjugated Alexa 488 (Thermo Fisher Scientific, D22910) injected at 0.2 mg/gm body weight. Tracers were made fresh in PBS, dissolved in 5 ul of PBS/gm body weight.

***In vivo* siRNA experiments**

250 nmol of Ambion's HPCL-IVR (HPLC grade, *in vivo* ready) pre-designed siRNAs targeting mouse vitronectin (Catalog No. 4457308, siRNA IDs s76001, s76002) and a control siRNA (Catalog No. 4457289) were ordered from ThermoFisher. Invivofectamine 3.0 (Thermo Fisher, IVF3001) was used to form siRNA complexes, as per manufacturer's instructions for systemic delivery into mice. siRNA duplexes were resuspended in DNase/RNase free water (Thermo Fisher, 10977) at 4.8 mg/ml, the complexation in Invivofectamine 3.0 was followed as per protocol. 100 ul of siRNA complex was injected into 6-week old mice (~20 gm) to result in 1 mg/kg dosing. siRNA complexes were injected into circulation through tail vein on 2 consecutive days. Subsequent experiments included confirmation of knock-down and leakage assays which were performed either 24 hours

(day 3) or 72 hours (day 5) post last siRNA injection. Confirmation of vitronectin knock-down was determined by ELISA on plasma isolated (see below) from mice. For leakage assays, Sulfo-NHS-biotin was injected (0.5 mg/gm body weight) into circulation via the tail-vein. Across all mice, retinas from left eyes were used for leakage analyses.

**Plasma isolation**

On the day of experiments, mice were briefly anaesthetized with 3% isoflurane and heparinized capillary tubes 1.1 mm x 7.5 mm (Thomas Scientific, 44B508) were used to collect blood from the retro-orbital sinus of the right eye. Collected blood was transferred to Eppendorf tubes containing 10 ul 0.5 M EDTA, pH 8.0 (Thermo Fisher, 15575020) and vortexed to prevent blood from coagulating. Isolated blood samples were centrifuged for 15 minutes at 2000xg at 4$^{o}$C and the supernatant was collected. Knockdown of vitronectin in plasma samples was confirmed by an ELISA kit for vitronectin (Molecular Innovations, MVNKT-TOT).

**Transmission electron microscopy**

HRP (Thermo Fisher Scientific, 31491), prepared fresh before each experiment was injected into the retro-orbital sinus of P10 pups briefly anaesthetized with isoflurane. HRP was injected at 0.5 mg/gm body weight, dissolved in 5 ul of PBS/body weight. Tracer was circulated for 10 minutes followed by enucleation of the contralateral eye and retina dissection in 4% PFA made in 0.1 M sodium cacodylate buffer (EMS, 11653). The brain was also dissected and both the tissues were first fixed in 5% glutaraldehyde (EMS, 16200) and 4% PFA mixture made in 0.1 M sodium cacodylate for 1 hour at RT. The brains and retinas were then fixed overnight at 4 $^{o}$C in 4% PFA in 0.1 M sodium cacodylate.

Following fixation both tissues were washed with 0.1 M sodium cacodylate buffer three times, 5 minutes each. 100 um sagittal sections of the brain and the whole retina were then incubated with fresh 0.5 mg/ml DAB (Sigma, D5905) for 25 min at RT. DAB was prepared in 0.1 M sodium cacodylate containing 0.05 M Tris-HCl and 0.01% hydrogen peroxide. Leaky areas within the retina and cerebellum were further microdissected, cut into 80 nm ultrathin sections and further processed for EM analysis as described previously (Chow and Gu, 2017).

**Western Blotting**

Retinas were dissected in PBS containing protease inhibitors and phosphatase inhibitor (Thermo Fisher Scientific, 87786 and 78420 respectively). Dissected retinas were further minced with a razor blade and allowed to lyse on ice for 30 minutes in lysis buffer (50 mM Tris-HCl, pH 7.4, 150 mM NaCl, 1% triton-X 100, 0.1% SDS supplemented with protease and phosphatase inhibitors). Lysed retinas were centrifuged for 15 minutes at 4$^{o}$C, the supernatant was collected and BCA assay (Thermo Fisher Scientific, 23225) was performed according to manufacturer's instructions to determine total protein concentrations. Samples were then analyzed for various proteins using SDS-PAGE and Western Blotting. The following antibodies were used: Pdgfrβ (Cell Signaling Technologies 3169, 1:500), Fibronectin (Abcam ab2413, 1:200)

### Cell culture media, growing conditions and coverslip coating

Mouse primary brain microvascular endothelial cells (Cell Biologics, C57-6023) were maintained in a complete mouse endothelial cell medium (Cell Biologics, M1168) supplied with VEGF, ECGS, Heparin, EGF, and FBS, according to the manufacture's instruction. To functionalize the surface of cover glasses, we first cleaned the cover glasses (Thomas Scientific, 1217N79) with air plasma (Harrick Plasma) for 15 min. For surface functionalization with single matrix proteins, the cleaned cover glasses were incubated with 50 µg/mL mouse collagen IV (Corning, 354233) in water, 50 µg/mL laminin 511 (Sigma-Aldrich, CC160) in PBS, or 50 µg/mL multimeric vitronectin (Molecular Innovations, HVN-U) in PBS, for 1 h at 37$^{o}$C. For surface functionalization with collagen IV plus laminin, the collagen IV-functionalized cover glasses were air dried and then incubated with 50 µg/mL laminin 511 in PBS for 1 h at 37$^{o}$C. For surface functionalization with Collagen IV plus laminin and vitronectin, the collagen IV-functionalized cover glasses were air dried and then incubated with 50 µg/mL laminin 511 and vitronectin in PBS for 1 h at 37$^{o}$C. The functionalized cover glasses were washed with cell culture media 3 times before cell plating. Cells were gently detached from tissue culture dishes using enzyme-free cell dissociation buffer (Gibco, 13151014) and then seeded on the functionalized cover glasses for at least 4 h before next treatment.

### Imaging focal adhesions in cell culture

To image integrin α5-mediated adhesion, the cells were snap chilled in ice-cold HHBSS, which is 1X HBSS (Gibco, 24020117) buffered with 10 mM HEPES (Gibco, 15630106). The cells were then incubated with primary antibody rat α-CD49e (α5, BD Biosciences 553319, 1:100) in HHBSS for 30 min at 4$^{o}$C and washed with ice-cold HHBSS 3 times for 5 min each. The cells were fixed in 4% ice-cold PFA in BPS for 15 min at RT and permeabilized with 0.1% triton-X in PBS for 15 min at RT. The cells were then washed with PBS 3 times for 5 min each and incubated in blocking buffer (5% bovine serum albumin in PBS) for 1 hour at RT. To simultaneously image paxillin, vinculin or pFAK with integrin α5, the blocked samples were incubated with corresponding primary antibody in blocking buffer for 2 h at RT and washed with PBS 3 times for 5 min each. Primary antibodies include mouse α-paxillin (BD Biosciences 610051, 1:100), mouse α-vinculin (Sigma-Aldrich V9131, 1:100) and rabbit α-FAK (phosphor Y397 from Abcam ab81298, 1:100). The samples were finally incubated with corresponding secondary antibodies at 1:500, phalloidin-iFluor 488 (Abcam, ab176753), and nuclear stain which is 1 µg/mL of either Hoechst 33342 (Thermo Scientific, 62249) or Ethidium Homodimer-1 (Thermo Scientific, E1169) in blocking buffer for 1 h at RT and washed with PBS 5 times for 5 min each before imaging.

### *In vitro* shRNA and endocytosis experiments

The DNA sequences encoding shRNA1 and 2 for integrin α5 knockdown are "CCCAGCAGGGAG-TCGTATTTACTCGAGTAAATACGACTCCCTGCTGGG" and "ATCAACTTGGAACCATAATTACTCGAGTA-ATTATGGTTCCAAGTTGAT", respectively. The DNA sequence encoding negative control or scramble shRNA is "CCTAAGGTTAAGTCGCCCTCGCTCGAGCGAGGGCGACTTAACCTTAGG", which is designed based on previously published scramble shRNA plasmid (from David Sabatini, addgene plasmid # 1864). These DNA fragments were cloned into pLKO.1 (a gift from David Root, Addgene plasmid # 10878) for RNA transcription in cells. The DNA encoding EBFP2 was cloned into the pLKO.1 vector in substitution of the sequence encoding neomycin resistant protein to report the transcription of shRNA. To package lentivirus for RNAi, HEK293FT cells (Invitrogen, R70007) in 35-mm tissue culture dishes were transfected with 1.5 µg pLKO.1 shRNA plasmid, 0.8 µg psPAX2 (a gift from D. Trono, Addgene plasmid # 12260), 0.7 µg pMD2.G (a gift from D. Trono, Addgene plasmid # 12259), and 9 µL Turbofect transfection reagent (Thermo Scientific, R0533) in 2 mL of Opti-MEM (Gibco, 31985062) for 12 h. The viruses were produced in DMEM supplied with 10 % FBS and 110 mg/mL sodium pyruvate (Gibco) and harvested 24 h after transfection. The mouse primary brain endothelial cells were infected with lentivirus and used for experiments 3 days after infection.

To test endocytosis, cells were washed with HHBSS 3 times and starved in HHBSS for 1 h at 37$^oC$. The cells were then incubated with 5 µg/mL of FM™ 1-43FX (Invitrogen, F35355) in HHBSS for 15 min at 37$^oC$ and then quickly washed with HHBSS 5 times. The cells were immediately fixed in 4% PFA in PBS for 15 min at RT and washed with PBS 3 times before imaging.

#### Imaging

Leakage assays in brain tissue were imaged on Olympus VS120 slide scanner. Rest of the fluorescent imaging was acquired on Leica TCS SP8 confocal. Z-stacks were obtained and all maximum-intensity projections are shown in all figures. TEM images were acquired on a 1200EX electron microscope (JOEL) equipped with a 2k CCD digital camera (AMT). All images were processed using FIJI. Cell culture imaging was acquired on an epi-fluorescence Leica DMI 6000B microscope.

### QUANTIFICATION AND STATISTICS

#### Fluorescent In situ hybridization

To determine percentage of pericytes positive for *Vtn* mRNA transcripts, PECAM1 immunostaining was first used to determine if cells positive for *Pdgfrb* transcripts were pericytes. Vessels containing only a single nucleus and ≤ 5 µm in width as shown in Figure 1C were identified as capillaries and *Pdgfrb*+ cells in close-contact or abutting the vessel were identified as pericytes (Figure 1C).

Scatter plots for *Pdgfrb* vs *Vtn* mRNA puncta was obtained by first identifying nuclei using the DAPI channel. DAPI containing nuclei were first segmented using FIJI and segmented regions were recorded in ROI manager. Fluorescent images of *Pdgfrb* and *Vtn* were thresholded using 'Otsu' method. *Pdgfrb* and *Vtn* puncta numbers in each of the segmented regions were obtained from the thresholded images using 'Find Maxima' with tolerance ≥ 30. Puncta numbers obtained for both transcripts in a given cell were plotted as X-Y scatter.

Scatter plot for *Itga5* or *Itgav* mRNA transcripts in endothelial cells vs pericytes was obtained in a similar manner. However, the DAPI nuclei segmented were first assigned as endothelial cells or pericytes using *Pecam1* and *Pdgfrb* mRNA respectively.

**Leakage assays**

Leakage was quantified as previously described (Andreone et al., 2017; Chow and Gu, 2017). For retinas, 770 µm x 770 µm areas across all 4 leaflets were first maximum intensity Z-projected, background subtracted, thresholded to obtain area of vessel (isolectin for retinas and ICAM2 for brain sections) and tracer (Sulfo-NHS-Biotin). 'Default' threshold method was used for thresholding isolectin signal while 'Li' thresholding was used for streptavidin (tracer) signal. Permeability index was defined as the ratio of area of tracer to area of vessel. Permeability index value of 1 implies tracer confined within vessels while values >1 indicate tracer extravasation from vessels. For leakage analysis of brain tissues, 6-8 sections were quantified per animal. Average of values across these regions/sections for a given animal was treated as one biological sample.

**ELISA for vitronectin protein expression levels**

ELISA analyses were performed using Arigo Biolaboratories free ELISA calculator at https://www.arigobio.com/elisa-analysis. Standard curve was generated by fitting the data to a 4-parameter logistic (4PL) curve. Our preliminary ELISA runs determined that plasma samples diluted 1:500 were usually in the range of the standard curve.

**Vesicular density from EM data**

For both retina and cerebellum, 15-20 vessels per animal were imaged under an electron microscope. Low magnification images encompassing complete blood vessel was first acquired using which the entire cytoplasmic area of each vessel was determined. For every blood vessel, tracer-filled vesicles were manually counted and then normalized to cytoplasmic area of the vessel (excluding area of nucleus) to obtain vesicular density. Average of vesicular density across all vessels of a given animal was considered one biological sample.

**Vessel density, branching, radial outgrowth**

Tilescan images to capture entire leaflets of retinas were acquired and these parameters were obtained for each leaflet. The average of all 4 leaflets was taken to be the measurement for a given animal. For vessel density measurements, isolectin stained vessel images were auto-thresholded to determine vessel area. Ratio of vessel area was normalized to total leaflet area to obtain vessel density measurements. For capillary branches, capillary branch points were counted manually in multiple 300 µm x 300 µm regions per leaflet, and the average across these regions was obtained for each animal. For P10 retinas, 300 µm x 300 µm regions were ideal as it allowed for the exclusion of arteries and veins. For radial outgrowth, vessel images were first auto-thresholded, distance from the optic nerve head to the tip of the retina was measured and this was normalized to distance from optic nerve head to edge of retinal tissue.

**Pericyte density and coverage**

Pericyte density and coverage analyses were performed as described previously (Chow and Gu, 2017). 770 µm x 770 µm areas across all 4 leaflets were first maximum intensity Z-projected and background subtracted. Erg and isolectin were auto-thresholded across all images and genotypes. For pericyte density measurements, DsRed signal was thresholded using 'Otsu' method to identify only pericyte cell bodies. The number of DsRed+ cell bodies was normalized to endothelial cell nuclei count to determine pericyte density. For pericyte coverage measurements, DsRed signal was thresholded using 'Li' method to allow detection of pericyte cell bodies as well as processes. Area of DsRed obtained was then normalized to area of thresholded isolectin to determine pericyte coverage.

**Astrocyte endfeet coverage from EM data**

Low magnification EM images were acquired to capture entire cross-section of vessels. The vascular basement membrane was first identified in each vessel (encompassing endothelial cells and pericytes) and perimeter of the cross-section was first obtained. Astrocyte endfeet coverage was defined as vessel perimeter in contact with astrocyte endfeet normalized to total perimeter of the vessel cross-section. Average of percent coverage of all vessels from one animal was reported as the percent coverage for that mouse.

**Western Blots**

All western blots were quantified using the gel analyzer tool in Fiji. Sample values were normalized to GAPDH values from corresponding lanes to account for differences in protein amounts across wells. The ratio of band intensity values of lysates from $Vtn^{-/-}$ mice to wildtype mice was then obtained to plot fold-change for mutant mice.

**Focal adhesions and dye uptake assays**

All images were processed using ImageJ Fiji. To identify integrin α5-mediated adhesions, the images were first background subtracted and thresholded. The particle analysis tool in Fiji was then used to quantify the number of adhesions in all imaging channels. The bright endocytic vesicles were identified by threshold tool and the ratio (%) of total fluorescence intensity in endocytic vesicles to the whole cell fluorescence intensity was measured to quantify endocytosis.

**Statistical Analysis**

All statistical analyses were performed using Prism 9 GraphPad software. For comparison between two groups, unpaired two-tailed Student's t-test was used. For comparison across multiple groups, one-way ANOVA with Tukey's post-hoc test was used.

**Data and code availability**

The codes used have been previously published and are freely available (see Methods description).

### Supplemental Figure Legends

**Figure S1. Vitronectin expression in CNS pericytes**

(A) Validation of vitronectin antibody (green) in retinas of P7 *Vtn*$^{-/-}$ mice. Vessels (isolectin) in magenta.

(B) In situ hybridization images showing *Vtn* (green), *Pdgfrb* (pericyte gene, magenta) and *Pecam1* (endothelial gene, red) in cortex and cerebellum of P7 mouse.

**Figure S2. Leakage seen in retinas of *Vtn*$^{-/-}$ mice is independent of tracer size and leakage persists through adulthood**

(A) Co-injection of Sulfo-NHS-Biotin (red) and 10 kDa Dextran (green) shows leakage (arrowheads) of both tracers from vessels (isolectin, blue) in P10 retinas of *Vtn*$^{-/-}$ mice.

(B) Quantification of 10 kDa Dextran leakage in wildtype and *Vtn*$^{-/-}$ mice. n = 5 animals per genotype. Mean ± S.D.; ***p < 0.001; Student's t-test.

(C) Leakage assay in 8-week old mice with Sulfo-NHS-Biotin (tracer, green) and vessels (isolectin, magenta). White boxes indicate the region shown in higher magnification images. Tracer-filled neuronal cell bodies (arrowheads) are indicated.

**Figure S3. Examples of vessels in cerebellum of *Vtn*$^{-/-}$ mice with basement membranes completely filled with HRP**

(C) EM Images of whole cross-sectional view of blood vessels in cerebellum of P10 mice. Lumen is filled with HRP. HRP-filled vesicles (arrowheads), pericytes (P) and nuclei outline (dashed lines) are indicated.

(D) Higher magnification images of boxed regions in (C) shows darker staining of basement membrane (black arrows) on the abluminal side (yellow dashed line) of *Vtn*$^{-/-}$ mice.

(D) Quantification of percentage of vessels with basement membrane filled with HRP in wildtype and *Vtn*$^{-/-}$ mice. n = 4 animals per genotype. Mean ± S.D.; ***p < 0.001; Student's t-test.

**Figure S4. Lack of vitronectin does not alter vascular basement membrane composition**

(A) Tilescan images of collagen IV in P10 retinas of wildtype and *Vtn*$^{-/-}$ mice.

(B) Higher magnification images of collagen IV (green), vessels (isolectin, blue) and pericytes (NG2:DsRed, red).

(C) Representative images of Collagen IV (green) ensheathment of vessels (isolectin, cyan) and pericytes (NG2:DsRed, red) in P10 retinas.

(D, E) Representative western blots (D) and quantification (E) of Fibronectin protein levels normalized to Gapdh in whole retinal lysates of P10 wildtype and *Vtn*$^{-/-}$ mice. n = 3 mice per genotype. Mean ± S.D.; n.s. not significant, p > 0.05; Student's t-test.

(F, G) Representative images of perlecan (F) and laminin α4 (G) in green and vessels (isolectin, magenta) in P10 retinas of wildtype and *Vtn*$^{-/-}$ mice. White boxes correspond to regions in the higher magnification images.

(H) Representative images of capillaries ensheathed by astrocyte endfeet. Cross-section of vessel and nucleus highlighted in red dashed lines, pericytes (P) shown in green, astrocyte endfeet (AE) in blue, L indicates vessel lumen filled with DAB.
(I) Quantification of percent of capillary cross-section covered by astrocyte endfeet in cerebellum of P10 wildtype and *Vtn*$^{-/-}$ mice. n = 4 mice per genotype, 15-20 vessels per animal. Mean ± S.D.; n.s. not significant, $p > 0.05$; Student's t-test.

**Figure S5. α5 containing adhesion structures are bonafide focal adhesions**
(A, B) Representative images (A) and quantification (B) of α5 adhesion structures in primary brain endothelial cells grown on collagen IV or laminin or vitronectin-coated dishes. n = 15 to 25 cells per condition from 2 independent experiments. Mean ± S.D.; n.s. not significant, $p > 0.05$, ***$p < 0.001$; one-way ANOVA with Tukey's post hoc test.
(C) Representative images of α5 containing adhesion structures (green) co-localizing with phospho-FAK (Y397) or paxillin or vinculin (red) in endothelial cells (phalloidin in magenta).
(D) Quantification of percentage of α5 adhesion structures positive for pFAK or paxillin or vinculin. n = 15 to 20 cells per co-staining from 2 independent experiments. In each case, >90% of α5 containing adhesions are positive for these focal adhesion markers.

**Figure S6. Endothelial-deletion of *Itga5* does not result in impaired vascular patterning or morphology**
(A) Tilescan images showing retinal vasculature of P10 retinas in mice lacking endothelial Itga5.
(B-D) Quantification of vessel density (B), capillary branching (C) and radial outgrowth (D) in retinas of wildtype and *Cdh5:CreER+; Itga5*$^{fl/fl}$ mice. n = 6 animals per genotype. Mean ± S.D.; n.s. not significant, $p > 0.05$; Student's t-test.

**Figure S7. Endothelial RGD-binding integrin, *Itgav*, is not required for CNS barrier integrity**
(A) In situ hybridization for *Itgav* (green) *Pecam1* (endothelial gene, red) and *Pdgfrb* (pericyte gene, magenta) in P7 brain tissue. See quantification in Figure 6B.
(B) Sulfo-NHS-Biotin (tracer, green) is confined to vessels (isolectin, magenta) in retinas of P10 *Cdh5:CreER+; Itgav*$^{fl/fl}$ mice. White boxes correspond to higher magnification images in (C).
(C) Tracer (green) confined to vessels with no leakage.
(D) Quantification of vessel permeability in wildtype and *Cdh5:CreER+; Itgav*$^{fl/fl}$ mice. n = 5 animals per genotype. Mean ± S.D.; n.s. not significant, $p > 0.05$; Student's t-test.
(E) Sulfo-NHS-Biotin (green) is confined to vessels (ICAM2, magenta) in the cerebellum of P10 *Cdh5:CreER+; Itgav*$^{fl/fl}$. White boxes correspond to higher magnification images depicting tracer (green) co-localizing with vessels (ICAM2, magenta).
(F) Quantification of vessel permeability in wildtype and *Cdh5:CreER+; Itgav*$^{fl/fl}$ mice. n = 5 animals per genotype. Mean ± S.D.; n.s. not significant, $p > 0.05$; Student's t-test.

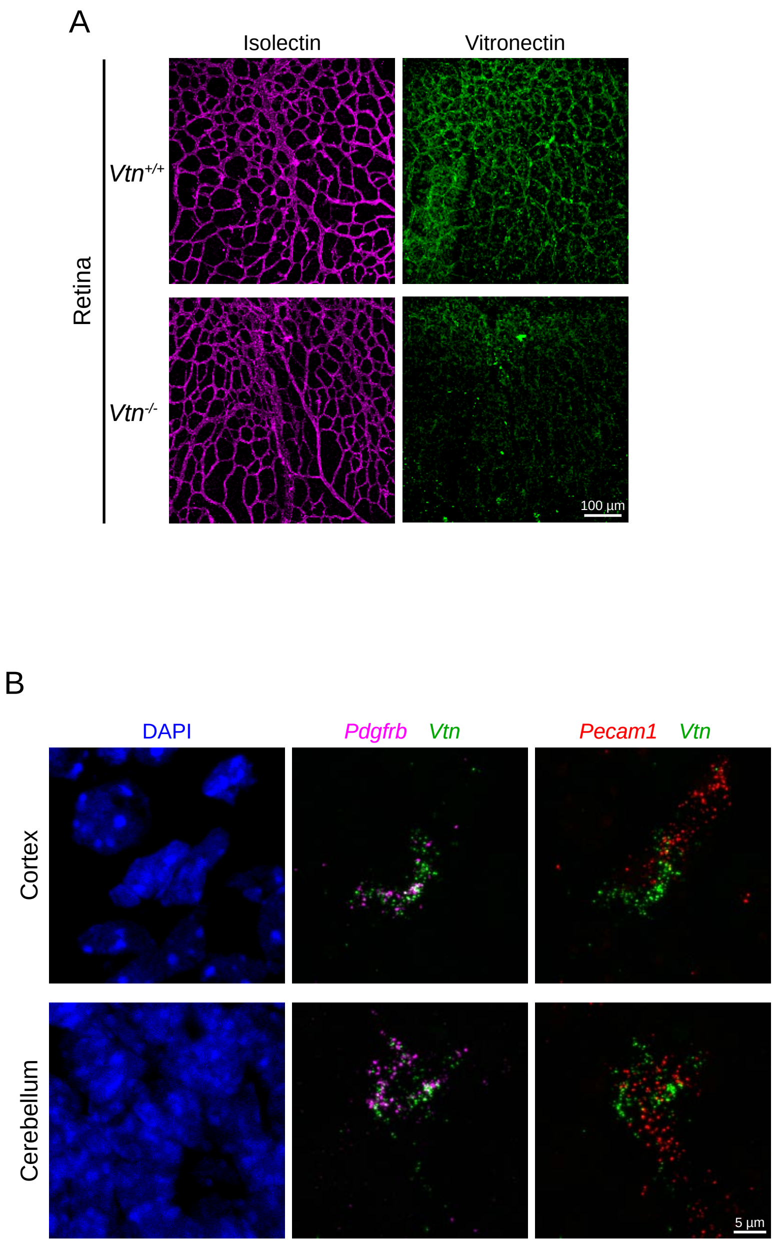


Ayloo et al. Supplementary Figure 1

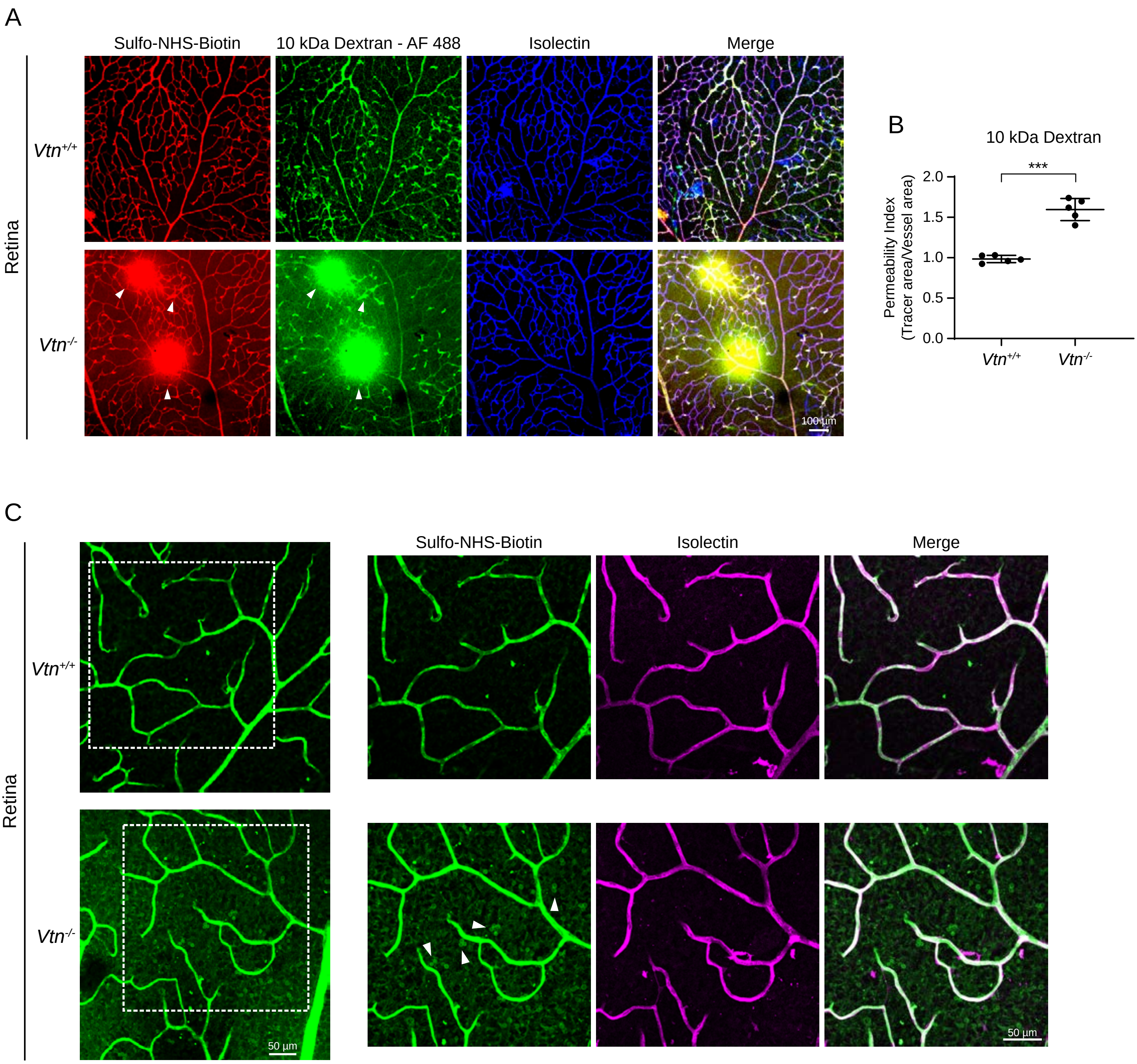


Ayloo et al. Supplementary Figure 2

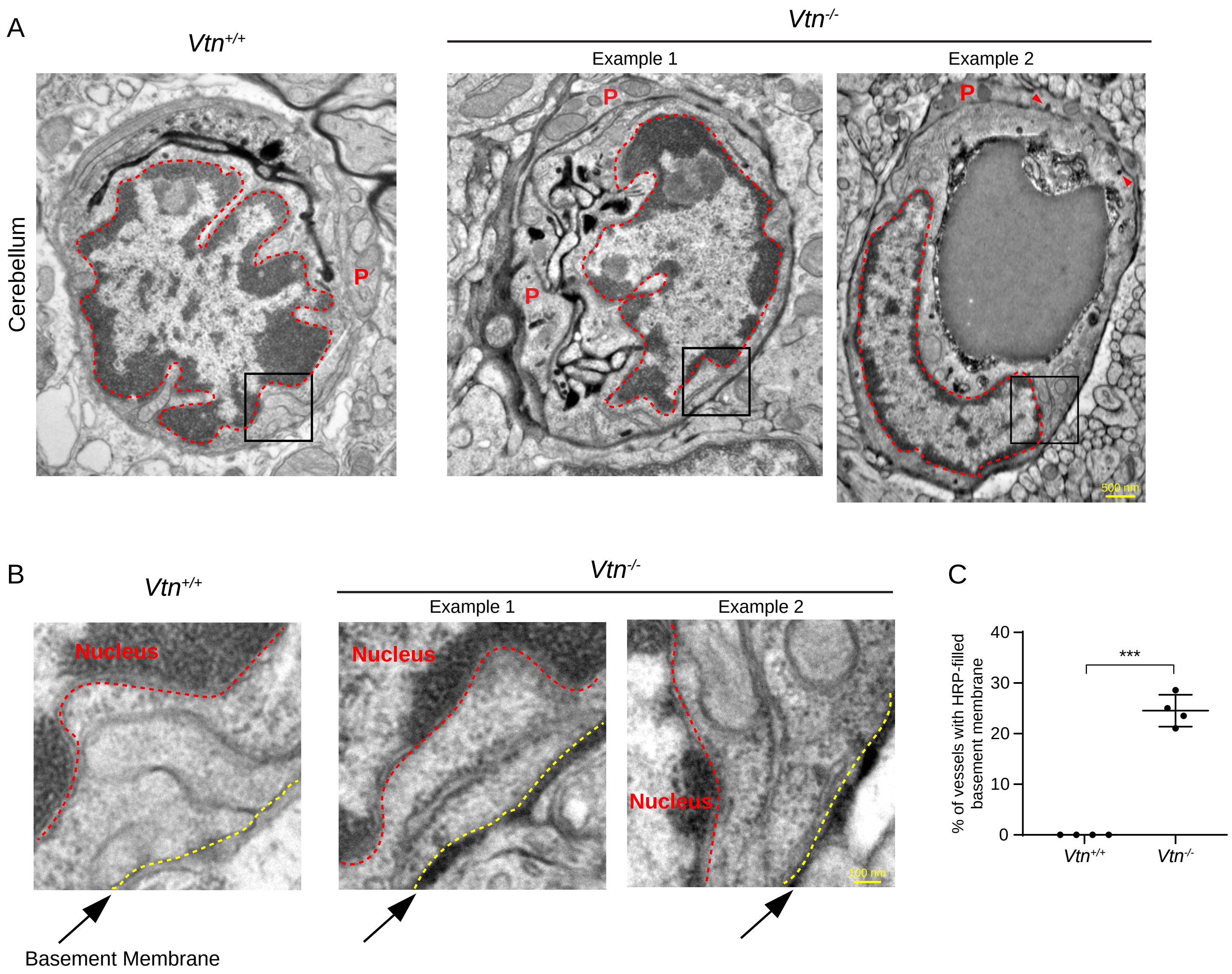


Ayloo et al. Supplementary Figure 3

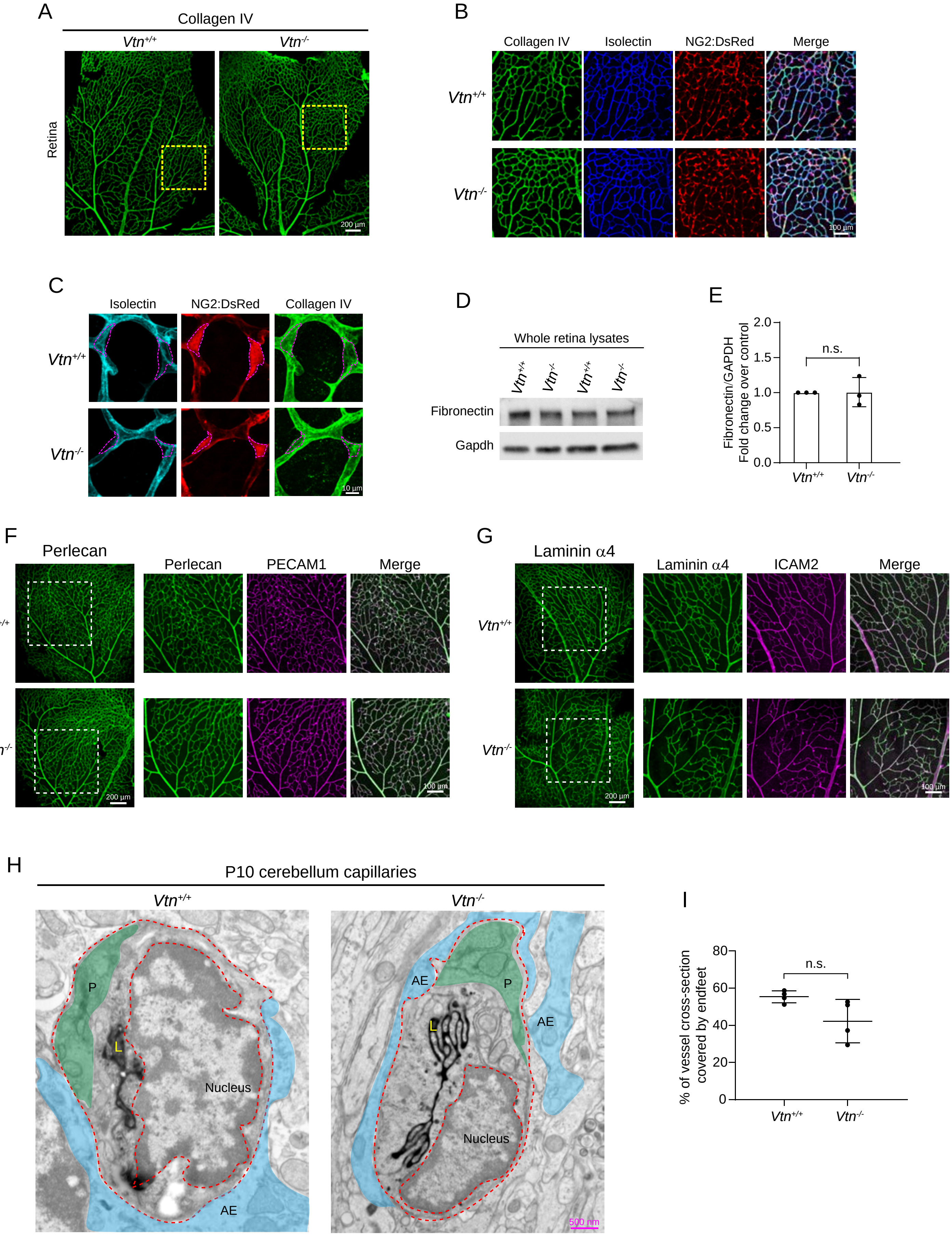


Ayloo et al. Supplementary Figure 4

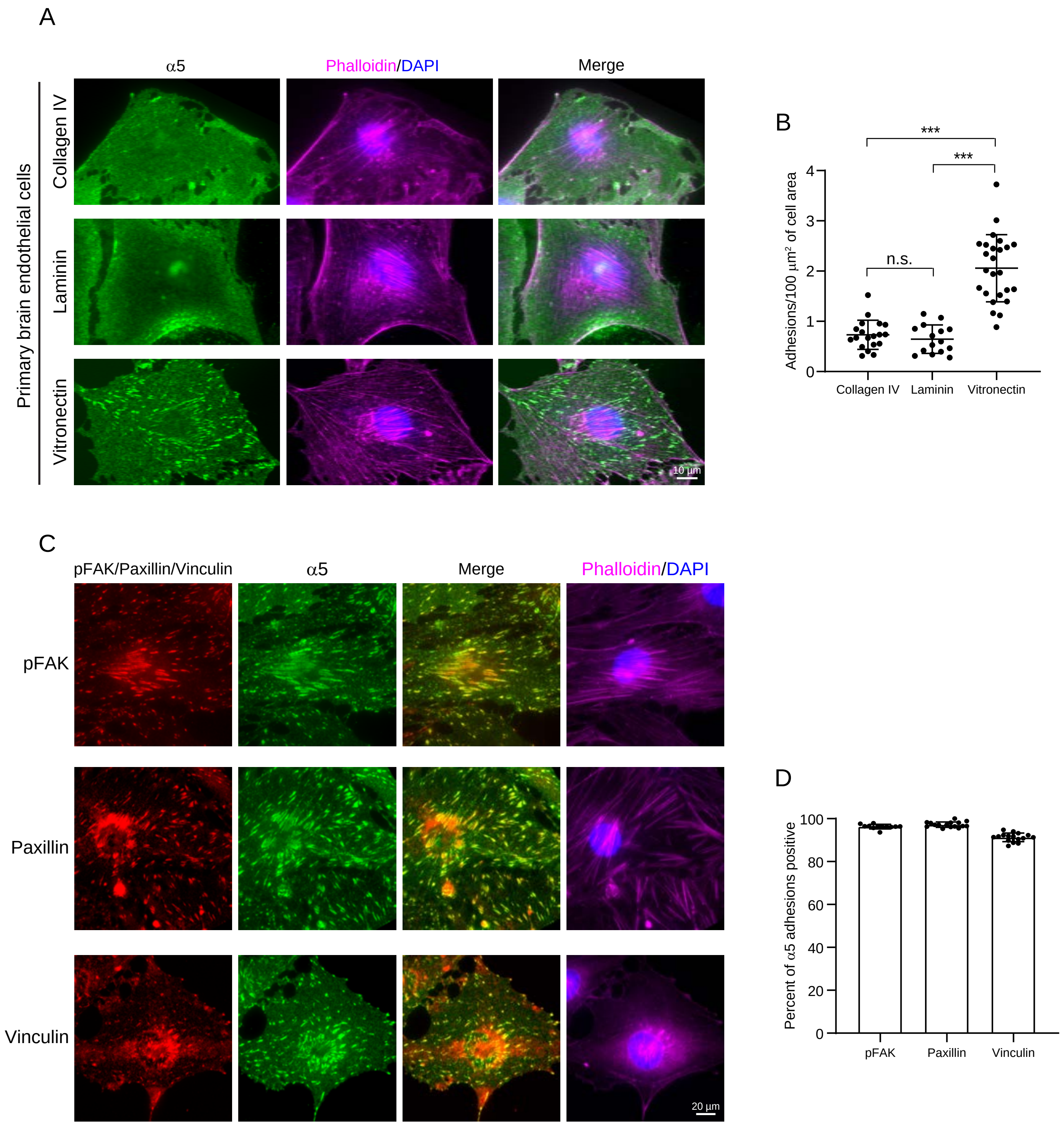


Ayloo et al. Supplementary Figure 5

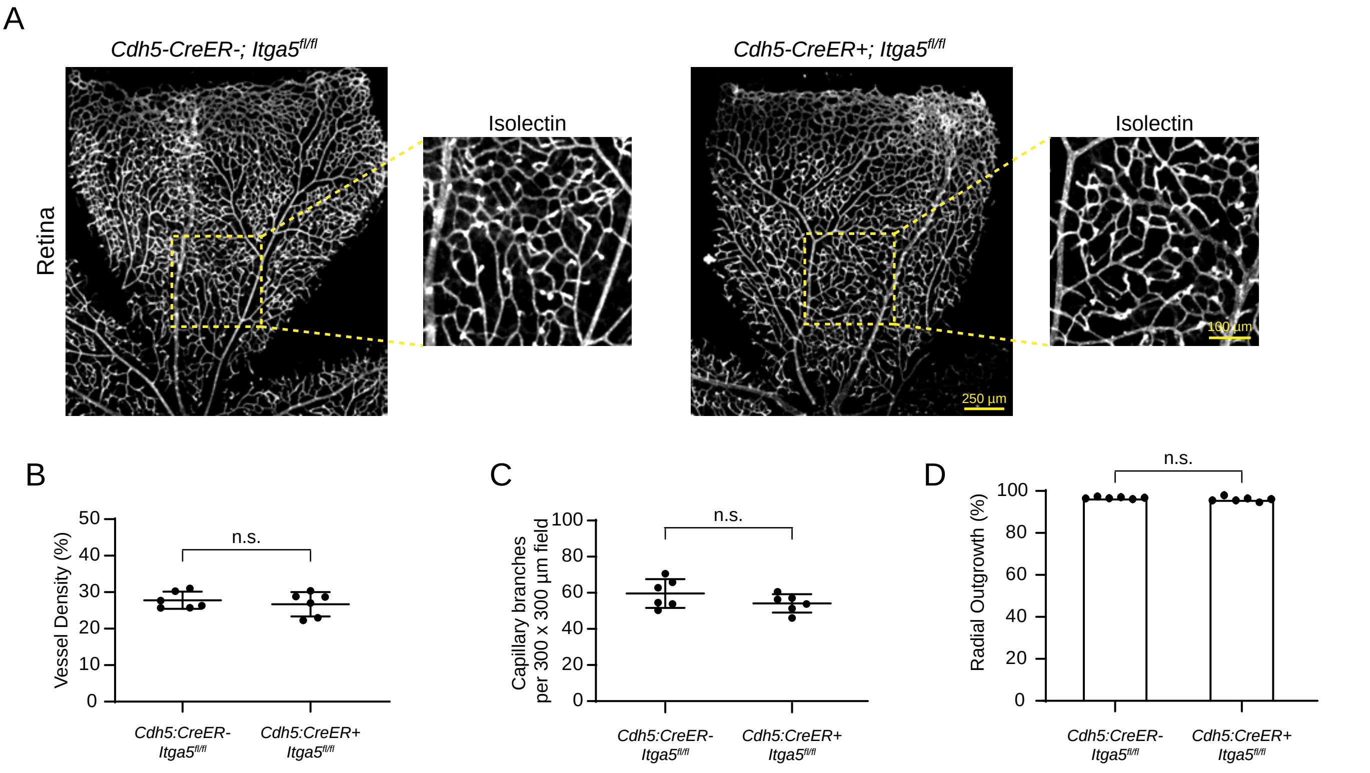


Ayloo et al. Supplementary Figure 6

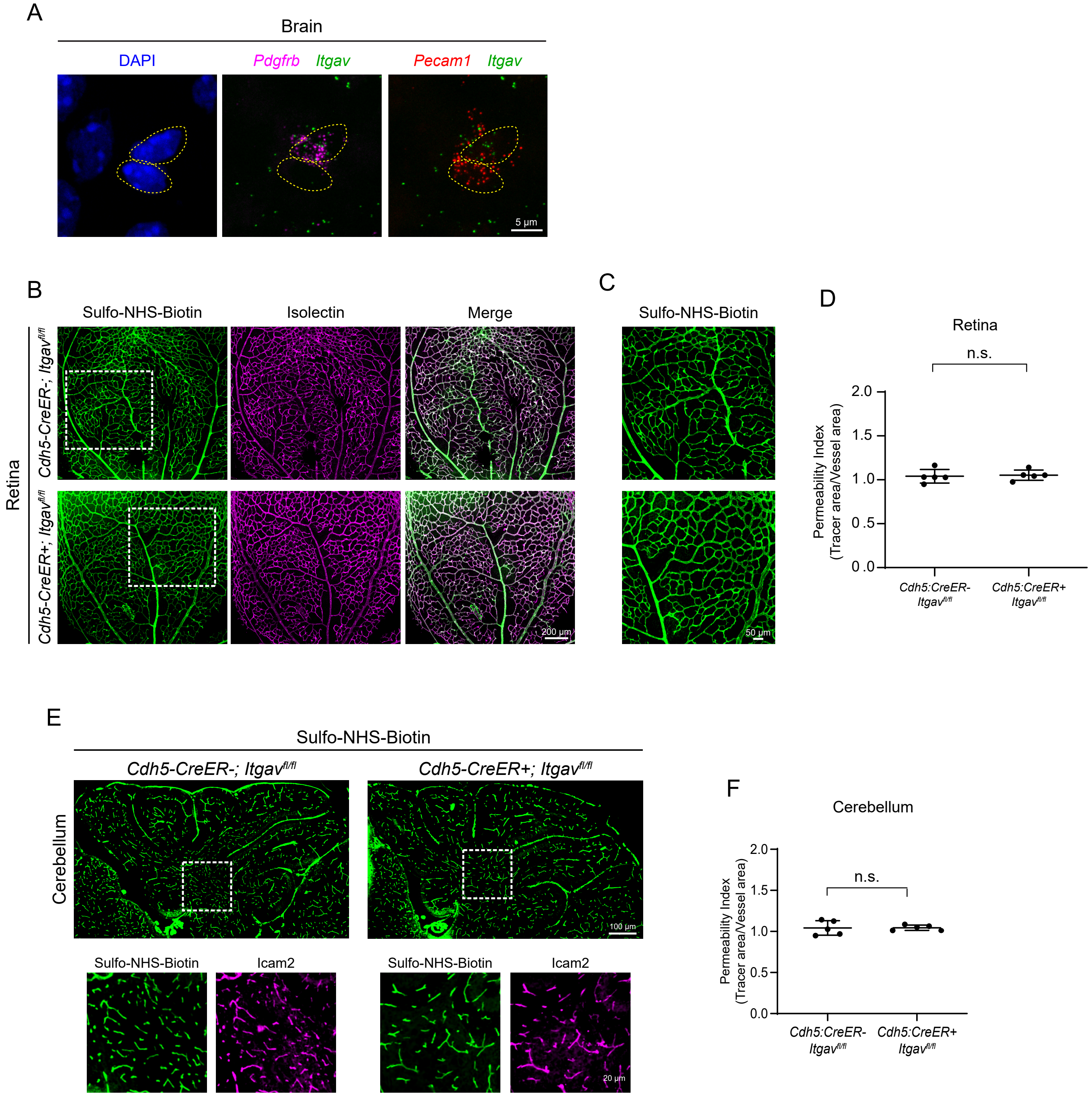


Ayloo et al. Supplementary Figure 7
